# Genome editing in the mouse brain with minimally immunogenic Cas9 RNPs

**DOI:** 10.1101/2023.04.13.536294

**Authors:** Elizabeth C. Stahl, Jennifer K. Sabo, Min Hyung Kang, Ryan Allen, Elizabeth Applegate, Shin Eui Kim, Yoonjin Kwon, Anmol Seth, Nicholas Lemus, Viviana Salinas-Rios, Katarzyna Soczek, Marena Trinidad, Linda T. Vo, Chris Jeans, Anna Wozniak, Timothy Morris, Athen Kimberlin, Thomas Foti, David F. Savage, Jennifer A. Doudna

**Affiliations:** Innovative Genomics Institute, University of California, Berkeley, California 94720, USA; California Institute for Quantitative Biosciences (QB3), University of California, Berkeley, California 94720, USA; Department of Molecular and Cell Biology, University of California, Berkeley, California 94720, USA; Aldevron LLC, Madison, Wisconsin 53719, USA; Howard Hughes Medical Institute, University of California, Berkeley, California 94720, USA; Department of Chemistry, University of California, Berkeley, CA, USA 94720; MBIB Division, Lawrence Berkeley National Laboratory, Berkeley, California 94720, USA; Gladstone Institutes, University of California, San Francisco, CA 94114

**Keywords:** CRISPR-Cas9, Genome Editing, Viral Vectors, Non-viral Delivery, Mouse, Brain, Host Immune Response, Neurons, Microglia, Endotoxin/LPS

## Abstract

Transient delivery of CRISPR-Cas9 ribonucleoproteins (RNPs) into the central nervous system (CNS) for therapeutic genome editing could avoid limitations of viral vector-based delivery including cargo capacity, immunogenicity, and cost. Here we tested the ability of cell penetrant Cas9 RNPs to edit the mouse striatum when introduced using a convection enhanced delivery system. These transient Cas9 RNPs showed comparable editing of neurons and reduced adaptive immune responses relative to one formulation of Cas9 delivered using AAV serotype 9. The production of ultra-low-endotoxin Cas9 protein manufactured at scale further improved innate immunity. We conclude that injection-based delivery of minimally immunogenic CRISPR genome editing RNPs into the CNS provides a valuable alternative to virus-mediated genome editing.

## Introduction

Editing somatic cells directly *in vivo* is anticipated to be the next wave of therapeutics for many genetic diseases, especially those affecting the central nervous system (CNS)^1,2^. Clustered regularly interspaced short palindromic repeats (CRISPR) is a revolutionary tool adapted from bacterial immune systems for genome editing^3–5^. To achieve gene disruption, the functional endonuclease, Cas9, is directed by a guide RNA to a target site in DNA to generate a double strand break leading to insertions and deletions (indels). Unfortunately, despite many genetic disease indications, the brain remains a challenging target for genome editing.

To circumvent the blood-brain barrier (BBB), most genomic medicines rely on direct intracranial injection of viral vectors encoding the transgene of interest. Viral vectors, such as recombinant adeno-associated virus (AAV), have had great success in gene therapy and are less immunogenic than most viral vectors, however, they require re-manufacturing for each target and are hindered by costly production scale up. Additionally, AAV has a limited DNA packaging capacity, and is associated with immunogenicity in the brain from both the vector and expression of foreign transgenes^6–9^. Although the brain has been considered an immune-privileged site, green fluorescent protein can induce a strong inflammatory response and neuronal cell death three weeks after injection with AAV serotype 9 has been reported^6–9^. Additionally, Cas9-specific immune responses have been elicited following AAV delivery in mice^10–12^ and pre-existing cellular and humoral immunity to Cas9 and AAVs are documented in humans^13–18^. Despite these drawbacks, AAVs are the most clinically relevant delivery systems currently in use for the CNS.

The development of transient, non-viral delivery systems that can effectively edit neurons throughout the brain with minimal immunogenicity would greatly facilitate future clinical applications. Previously, we developed cell penetrating Cas9 ribonucleoproteins (RNPs) capable of genome editing in mouse neurons both *in vitro* and *in vivo*^19^. To enable self-delivery of the Cas9 RNPs, four repeats of the positively charged Simian vacuolating virus 40 nuclear localization sequences (SV40-NLS) were fused to the N-terminus along with two repeats to the C-terminus of Cas9, a strategy that was also reported to enable delivery of zinc-finger nucleases^20^. Using a single guide to turn on the tdTomato reporter from the lox-stop-lox (LSL-Ai9^21^) mouse, we reported edited striatal volume of approximately 1.5mm^3^ ^19^.

Here we report further optimization of cell penetrant Cas9 RNPs, demonstrating efficacy in human primary cells and improved editing of the mouse striatum using a convection enhanced delivery (CED) system. We compared the transient RNP complexes to AAV serotype 9 for Cas9 delivery to the CNS, to measure both editing efficiency and the host immune response. We found that the Cas9-AAV was able to better diffuse throughout the brain, leading to distally edited cells; while the cell penetrant Cas9-RNPs edited significantly more neurons within the region near the injection site. Both groups elicited humoral responses, but vehicle-specific antibodies in the Cas9-AAV group persisted at high levels out to 90-days. Cas9-AAV treated brains were also associated with significantly elevated *Cd3e* gene expression at four weeks, suggesting an ongoing adaptive immune response. Cas9-RNP treated brains showed acute microglial activation that was mitigated by reducing endotoxin levels during protein manufacturing scale up. Taken together, Cas9 RNPs are a promising strategy for future therapeutic intervention in neurological disorders to address current limitations of viral delivery.

## Results

### Development of Cas9 cell penetrant RNP and AAV to measure genome editing with the tdTomato reporter system

Creating a large deletion in the lox-stop-lox cassette in Ai9 mice with a single guide RNA (sgRNA) enables expression of tdTomato and efficient quantification of editing by fluorescent read out (Figure S1A). Cas9 from *Streptococcus pyogenes* (engineered with four copies of SV40 NLS on the N-terminus and two copies on the C-terminus (4x-SpyCas9-2x) to be cell penetrant) was first produced from recombinant *E. coli* in a laboratory setting, using a low-endotoxin method. Editing efficiency of the RNP was compared to Cas9 delivered by recombinant adeno-associated virus (AAV) (Figure 1A-B). Since SpyCas9 cannot be packaged within a single AAV with its guide RNA, we used clinically relevant AAV-SauCas9-sgRNA (derived from *Staphylococcus aureus*)^22–24^. AAV serotype 9 was produced using a baculovirus transfected into Sf9 insect cells^25,26^. To control for differences in the Cas9 orthologs, cell penetrant 4x-SauCas9-2x protein was also produced following the same expression and purification methods as 4x-SpyCas9-2x. Due to differences in PAM requirements between the two Cas9 orthologs (SpyCas9 NGG, SauCas9 NNGRRT), a new guide was designed for SauCas9 to target the tdTomato locus (Figure S1B).

**Figure 1.**
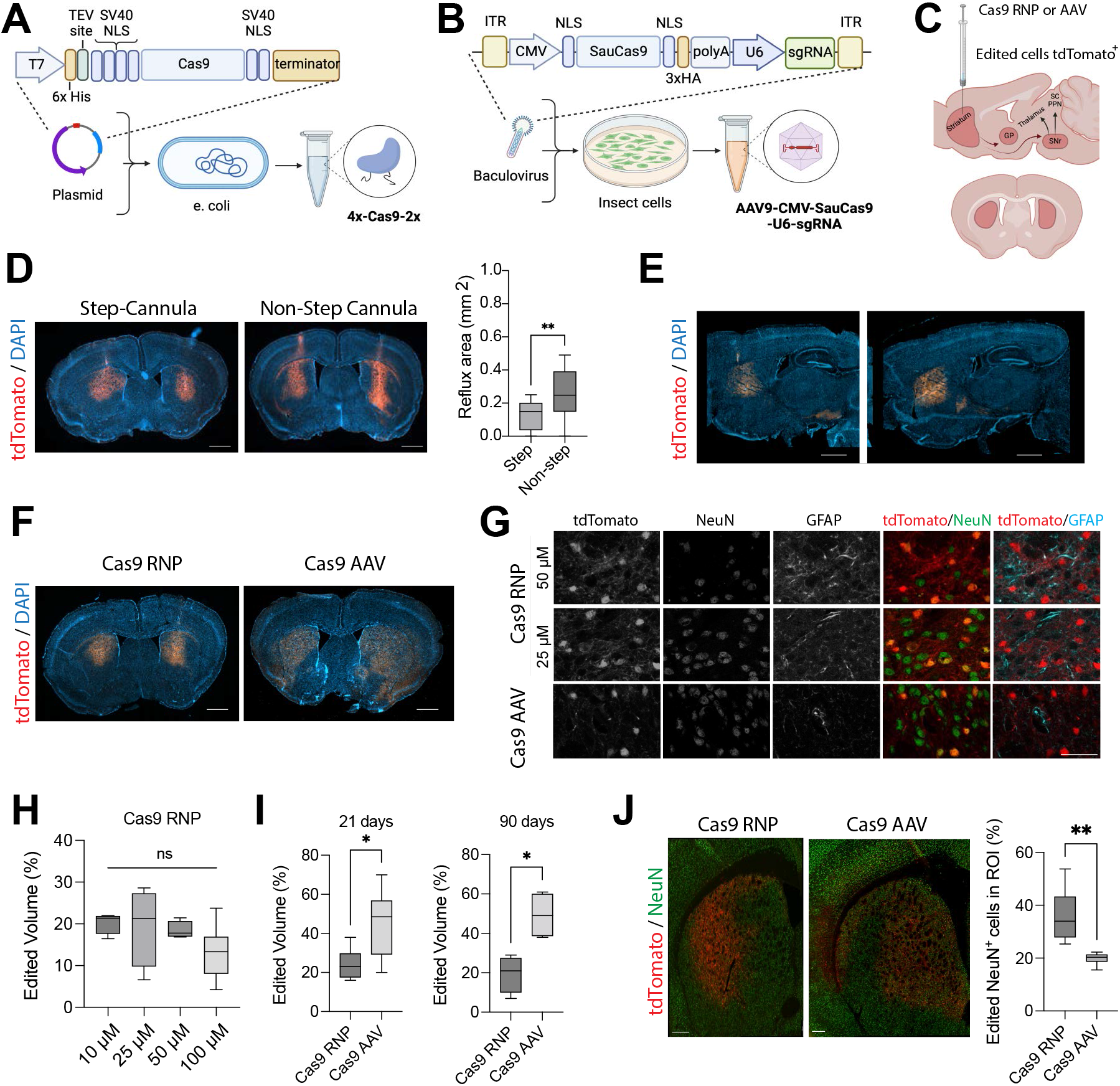
In vivo editing at tdTomato locus with viral and non-viral Cas9 delivery strategies. (A) Schematic of 4x-SpyCas9-2x cell-penetrating protein expression and purification systems, (B) AAV9-SauCas9-sgRNA expression and purification systems, (C) and expected edited brain regions in the partial basal ganglia direct circuit shown in sagittal view (top) and coronal view (bottom). Neurons extend from the striatum into the globus pallidus (GP) and substantia nigra (SNr). Created with BioRender.com. (D) Comparison of convection enhanced delivery (CED) of cell-penetrant 4x-SpyCas9-2x RNP with step and non-step cannulas. The step-cannula significantly reduced reflux of material in the needle injection track (n=3-6 injections per group, unpaired t-test, ** p<0.01.) Scale bar: 1 mm. (E) Serial sections of single hemisphere sagittal view of edited tdTomato+ cells in the basal ganglia circuit after injection of Cas9 RNP with CED into the striatum, with signal detected near the GP and SNr. Scale bar: 1 mm. (F) Representative coronal section of the striatum of mice that received Cas9 RNPs and AAVs at 21 days post-injection, showing the distribution of tdTomato+ edited cells. Scale bar: 1 mm. (G) Co-staining of tdTomato with NeuN and GFAP in the striatum at 90-days post-injection. Scale bar: 50 µm. (H) Volume of edited striatal tissue as the concentration of injected Cas9 RNPs was increased from 10 to 100 µM (n=4-6 injections, one-way ANOVA, ns). (I) Quantification of editing following treatment with Cas9 AAV (3e9 vg/ µL, 1.5e10 vg/hemisphere) and Cas9 RNPs (25 µM, 125pmol/hemisphere) at 21 and 90-days (n=4-6 injections, one-way ANOVA, * p< 0.05). (J) Co-localization of tdTomato and NeuN quantified per regions of interest (ROI), e.g., edited area per hemisphere (n=4-6 injections, one-way ANOVA, ** p< 0.01). Scale bar: 250 µm.

We confirmed editing in neural precursor cells (NPCs) isolated from embryonic day 13.5 Ai9 mice with all constructs *in vitro* (Figure S1). Addition of 4x-SauCas9-2x RNP into the cell culture supernatant enabled editing of NPCs compared to 0x-SauCas9-2x RNPs, demonstrating that the four SV40-NLS peptides can mediate delivery of additional Cas9 orthologs. 4x-SauCas9-2x RNPs were slightly less efficacious than 4x-SpyCas9-2x RNPs, which could be explained in part by differences in the guide RNAs (Figure S1A-B). We observed editing of mouse NPCs with SauCas9 when delivered as both cell penetrant RNP (59 ± 6% tdTomato cells^+^, Figure S1D) and AAV (42 ± 1% tdTomato cells^+^, Figure S1E). The same titer of AAV9-CMV-GFP resulted in 96 ± 2% GFP^+^ cells, suggesting that editing lagged behind transduction; and an increase in empty capsids in the AAV-CMV-SauCas9 group was noted (Figure S1F)^27,28^.

To further examine the potential for cell penetrant Cas9 RNPs to edit difficult cells *in vitro*, we tested delivery and editing with 4x-SpyCas9-2x in human neural precursor cells derived from induced pluripotent stem cells (iPSCs)^29–31^. Human NPCs were treated with pre-formed RNPs using an established guide RNA targeting EMX1^32^. In human cells, we detected 10-fold higher rates of editing with 4x-SpyCas9-2x compared to standard RNPs (0x-SpyCas9-2x) delivered with commercial transfection reagents (Figure S2A-B).

### Cas9 RNPs result in modest editing of brain parenchyma following delivery into cerebrospinal fluid

To determine the optimal route of delivery for Cas9 RNPs into the mouse CNS, we tested intraparenchymal injections into the striatum, as well as injection into the cerebrospinal fluid (CSF), including intrathecal (IT) and intracerebroventricular (ICV) routes. Following IT injection of cell penetrant RNPs, we observed edited glial cells and neurons in the cortex and striatum of one hemisphere, but no editing within the spinal cord (Figure S3A). Following ICV injection of Cas9 RNPs in neonatal p0 mice, we observed tdTomato^+^ cells in the subventricular zone and white matter, including glial cells and neural stem/progenitor cells expressing Ki67 and DCX evaluated three weeks after delivery (Figure S3B-C). Editing post-ICV injection in adult mice was restricted to the cells within the lateral ventricles, choroid plexus, subventricular zone, and hippocampus in a subset of mice (Figure S3D-E). The total number of edited cells with RNP delivery into the CSF was lower than with direct intraparenchymal injection (Figure S3F).

Therefore, we sought to further improve upon intraparenchymal injections using a convection enhanced delivery system (CED), which generates a high-pressure gradient to aid in biodistribution of macromolecules in the brain. CED has been used to increase the spread of AAV in the brains of large animal models and humans by infusing relatively high injection volumes at high rates^33,34^. Additionally, we tested two needle designs to enable CED with Cas9-RNPs. We found that the CED enabled robust editing in the mouse striatum (Figure S4B) and the step-cannula reduced reflux of RNP from the needle-injection track, as reported previously^35^ (Figure 1D). Furthermore, tdTomato^+^ neurons edited by Cas9-RNPs within the striatum were observed to extend along the basal ganglia circuit into the globus pallidus and substantia nigra along the anterior-posterior axis (Figure 1C-E).

### Convection enhanced delivery of Cas9 RNPs and AAVs mediates robust editing in the mouse striatum

Using bilateral CED injections into the striatum, we compared edited tissue volume with the 4x-SpyCas9-2x RNP, 4x-SauCas9-2x RNP, and AAV9-SauCas9-sgRNA in adult Ai9 mice at three weeks post-injection. Despite performing well *in vitro*, 4x-SauCas9-2x RNPs underperformed *in vivo* when tested at two different doses and additional NLS configurations (Figure S5A-G, S6D). Therefore, we performed our primary comparison with two orthologous systems: 4x-SpyCas9-2x RNP (hereafter referred to as Cas9-RNP) and AAV9-SauCas9-sgRNA (hereafter referred to as Cas9-AAV, which serves as a positive control, Figure 1F).

First, we tested Cas9-AAV injection with CED at two doses, 3×10^8^ vg/µL and 3×10^9^ vg/µL (1.5×10^9^-1.5×10^10^ vg/hemisphere)^36^, and proceeded with the higher dose for subsequent studies *in vivo* (Figure S6A-B). We also tested several doses of Cas9-RNP ranging from 10µM to 100µM (50-500 pmol/hemisphere). Interestingly, there was no significant difference in editing when delivering RNPs in this concentration range (Figure 1H, Figure S6C). We chose the 25µM RNP concentration (4.15 mg/mL or approximately 1.75 mg/kg Cas9) group for further study as it had the highest maximal editing rate. Above 25µM in the RNP group, we observed a decrease in NeuN staining and an increase in GFAP staining out to 90-days in the Cas9-RNP group, suggesting dose-limiting effects (Figure 1G).

At both 21 and 90-days post-injection, the Cas9-AAV group outperformed the Cas9-RNP group when quantifying total edited striatal volume (n=8 at 21-days, n=4 at 90-days, p<0.05, Figure 1I). The volume of edited cells was relatively stable in the Cas9-AAV group between 21 and 90-days at approximately 47 ± 3% (covering approximately 13.4 mm^3^ of striatum), while the Cas9-RNP group had editing levels of 22 ± 3% (approximately 6.2 mm^3^ of striatum) between 21 and 90-days (increased from previous report of editing 1.5mm^3^ striatal volume^19^). Edited cells were observed further along the rostral-caudal axis in the Cas9-AAV group (-2.12 mm to 2.5 mm relative to Bregma), demonstrating better diffusion of the editor away from the injection site (Figure S6E-F).

Since large deletions in the tdTomato locus make on-target editing difficult to assess using short-read next-generation sequencing (NGS), we developed an NHEJ droplet digital PCR assay (ddPCR) to measure drop-off of HEX-labeled probes over the cut sites, in relation to distal reference FAM-labeled probes. Genomic DNA was isolated from 2-mm thick sections of each injected hemisphere, covering multiple brain sub-structures. Loss of the HEX probe reached 2 ± 1% in the Cas9-RNP group and 15 ± 10% in the Cas9-AAV group, indicating edited alleles, when measured at 28-days (Figure S7).

We also quantified the percentage of edited NeuN^+^ neurons within the tdTomato^+^ region of interest (ROI) per hemisphere between Cas9-AAV and Cas9-RNP at the 21-day timepoint. We found that Cas9-RNP edited significantly more NeuN^+^ neurons per ROI (36 ± 10%) compared to Cas9-AAV (20 ± 2%) (Figure 1J, n=6-8 injections, p<0.05). Within the ROI, neurons were the most frequently edited cell type in both groups, including DARPP-32^+^ medium spiny neurons (Figure S8). Additionally editing of ALDH1L1^+^ and OLIG2^+^ glial cells was noted in both groups (approximately 2% of edited cells within the ROI in the Cas9-RNP group and 8% of cells in the Cas9-AAV group). Therefore, Cas9-RNPs were able to edit comparable numbers of neurons and glia as Cas9-AAVs in a given area of striatal tissue.

### Comparison of local and peripheral immune response between Cas9 RNPs and AAVs in the Ai9 reporter mouse

We next examined the local and peripheral immune response following delivery of Cas9 RNPs and AAVs into the brain. Using immunofluorescent staining for Iba1 (Figure 2A), we observed a significant increase in percent Iba1^+^ area in the 25µM Cas9-RNP group from sham-treated animals (Figure 2B, n=6 replicates, p<0.05). Staining for CD45 showed dim expression on Iba1^+^ microglia and high expression on CD3^+^ T-cells, which were slightly increased in the 25µM Cas9-RNP group compared to the sham and Cas9-AAV, but not significantly different, at three weeks post-injection (Figure 2C-D, n=6-12 replicates).

**Figure 2.**
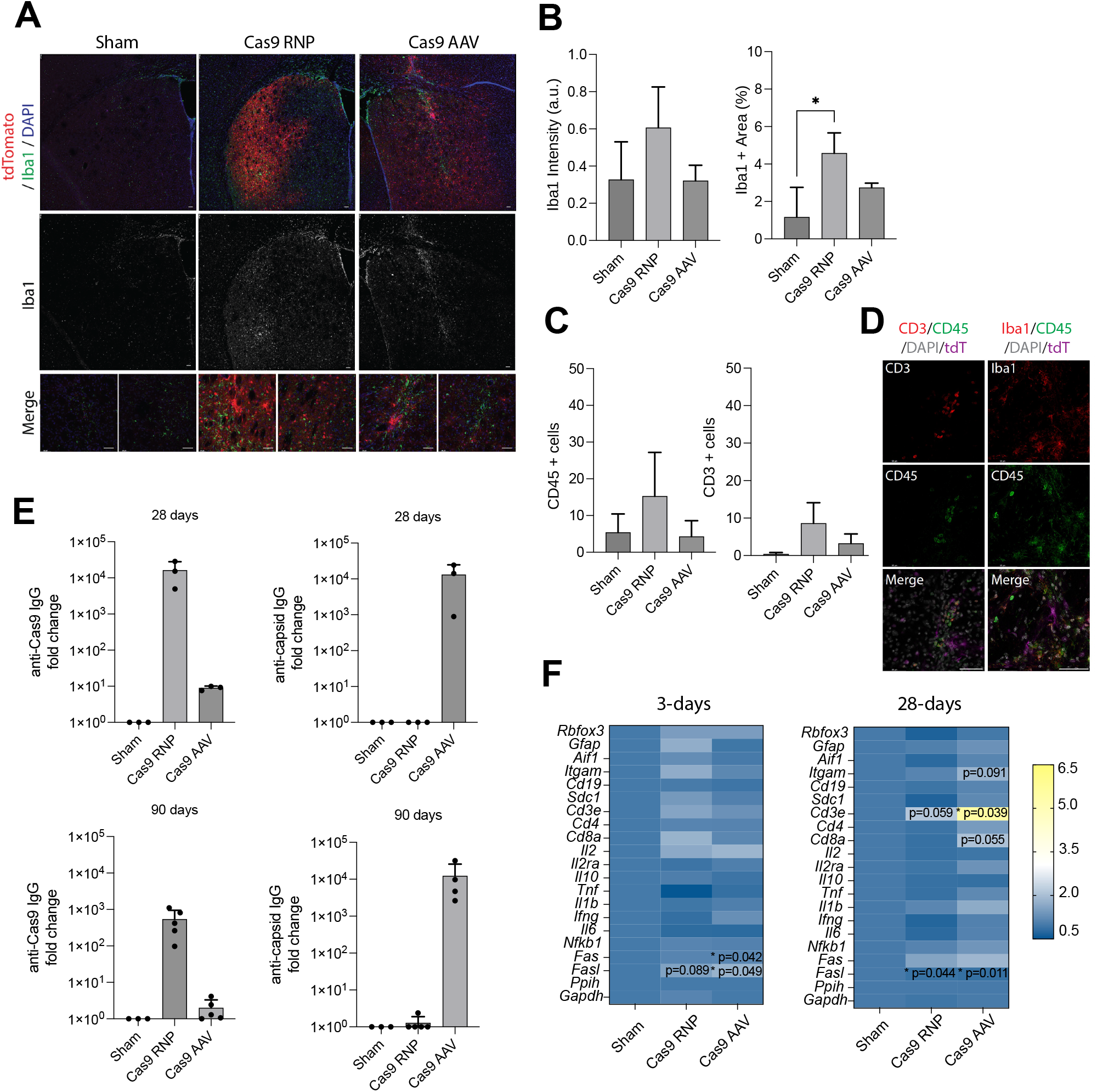
Immune response following in vivo editing with viral and non-viral Cas9 delivery strategies. (A) Representative immunostaining of Iba1 (microglia, green) with tdTomato and DAPI using confocal microscopy. Scale bar: 50 µm. (B) Quantification of Iba1^+^ staining intensity and percent area (n=4-6 technical replicates, one-way ANOVA, *p<0.05). (C) Quantification of CD45^+^ and CD3^+^ cells per image (n=3-6 replicates, one-way ANOVA, ns). (D) Representative images of CD45, CD3, and Iba1 showing co-expression of CD45 (green) with both Iba1 (microglia, red) and CD3 (T-cells, red) cells and differential cell morphology. Merged images include DAPI (gray) and tdTomato (magenta). Scale bar: 50 µm. (E) Quantification of IgG antibodies against Cas9 or AAV capsid proteins measured 28 and 90-days after bilateral intrastriatal injections by ELISA (n=3-5 biological replicates). (F) Heat map summarizing RT-qPCR results of gene expression from homogenized brain tissue near the injection site (striatum and cortex) at two time-points*. Ppih* was used as a housekeeping control for delta-delta Ct analysis and compared to the sham group using Qiagen analysis portal (n=4, *p < 0.05).

In addition to the immune response at the local site of injection, circulating IgG antibodies were measured at 28 and 90-days post-injection. We found that sham-treated animals had no pre-existing antibodies to either SpyCas9, SauCas9, nor AAV9 capsids. At 28-days following striatal injection, there was a 1.6e4-fold increase in anti-SpyCas9 IgG in the 25µM Cas9-RNP group, a 1.3e4-fold increase in anti-AAV9 capsid IgG in the Cas9-AAV group, and an 8.9e1-fold increase in anti-SauCas9 IgG in the Cas9-AAV group (i.e., humoral response against transgene) (Figure 2E, n=3-5 biological replicates). No cross-reactivity was observed between ortholog RNPs, as described previously^11^, nor were any anti-AAV capsid antibodies detected in the RNP group. At 90-days, the levels of IgG fell to a 5.4e2-fold increase in the 25µM Cas9 RNP group and 1.2e4-fold increase in the Cas9 AAV group from the sham controls, demonstrating greater maintenance of systemic antibodies against the capsid in the AAV group. The cellular and humoral immune response to Cas9 RNPs was dose-dependent and a significant increase in CD45^+^ cells was observed at the 100µM RNP dose compared to sham, Cas9-AAV, and Cas9-RNP at 25µM (Figure S9). Cas9-reactive cells were also identified in the spleen by interferon-gamma (IFN-*γ*) ELISpot assay (Figure S9E) at both 25µM and 100µM doses of Cas9-RNPs, but not in sham treated animals.

Since the mice had no pre-existing antibodies to SpyCas9, we tested how the immune response would differ in the RNP group by first exposing the mice to a single subcutaneous injection of 4x-SpyCas9-2x protein and adjuvant (AddaVax^TM^) four-weeks prior to stereotaxic surgery with Cas9-RNPs. We found that pre-exposing the mice to Cas9 with adjuvant had a synergistic effect on both serum IgG and activation of IFN-*γ*^+^ cells in the spleen (Figure S9F-I). Mice that received surgery maintained tdTomato^+^ cells in the brain to the measured time point. Additional studies using this immunization strategy may help to further characterize the immune response to Cas9-RNPs by modeling pre-existing immunity relevant to humans.

Finally, we measured gene expression changes near the injection site in mice that received Cas9-RNP and AAV at 3 and 28-days post-injection using RT-qPCR. At three days, the Cas9-AAV group had a modest but significant increase in *Fas* (1.19-fold) and *Fasl* (1.85-fold) compared to the sham group (Figure 2F). At 28-days post-injection, both Cas9-RNP and - AAV had a significant increase in *Fas* (1.61 and 1.89-fold respectively). In addition, the Cas9-AAV group had a significant increase in *CD3e* gene expression (5.45-fold, n=4 replicates, p<0.05), closely followed by *CD8a* (2.06-fold, ns, p=0.06), while Cas9-RNP had a slight but non-significant increase in *CD3e* (2.15-fold, ns, p=0.06), compared to the sham group.

There were no detectable off-target editing events at 1 and 4-months post-injection in any of the experimental groups at the evaluated sites (Figure S10A-C). In the Cas9-AAV group, the Cas9 transgene was expressed in the brain out to 4-months, the last tested time point, as expected (Figure S10D). Additionally, few genes were differentially expressed between the groups at 4-months, except for *Fas* (1.54-fold, p<0.05), which was significantly elevated in the Cas9-AAV group compared to the sham (Figure S10E-F). We used long read sequencing to examine whether any fragments of the viral genome had been integrated near the cut site in the tdTomato locus, as previously reported^37–40^. We also observed partial integrations of viral fragments in our amplicon, although our *in vivo* editing rates and sequencing depth were relatively low (Figure S11).

Overall, delivery of Cas9 by either AAV or RNP resulted in significant activation of immune responses in the brain and periphery, although with generally small effect sizes compared to sham injected mice. The increased Iba1^+^ cells near tdTomato^+^ cells in the striatum of the 25µM Cas9-RNP group raised the question of whether the response was due to Cas9 itself or impurities within the protein product. We hypothesized that the local immune response may be due to endotoxins from *E. coli* in the RNP complexes.

### Production and testing of ultra-low endotoxin 4x-SpyCas9-2x protein

To examine the impact of endotoxin on the immune response to RNPs, we partnered with a commercial producer of Cas9 protein and were able to significantly scale up manufacturing to produce a large quantity of ultra-low endotoxin 4x-SpyCas9-2x protein using an industrial tag-free expression and purification system (Figure 3A).

**Figure 3.**
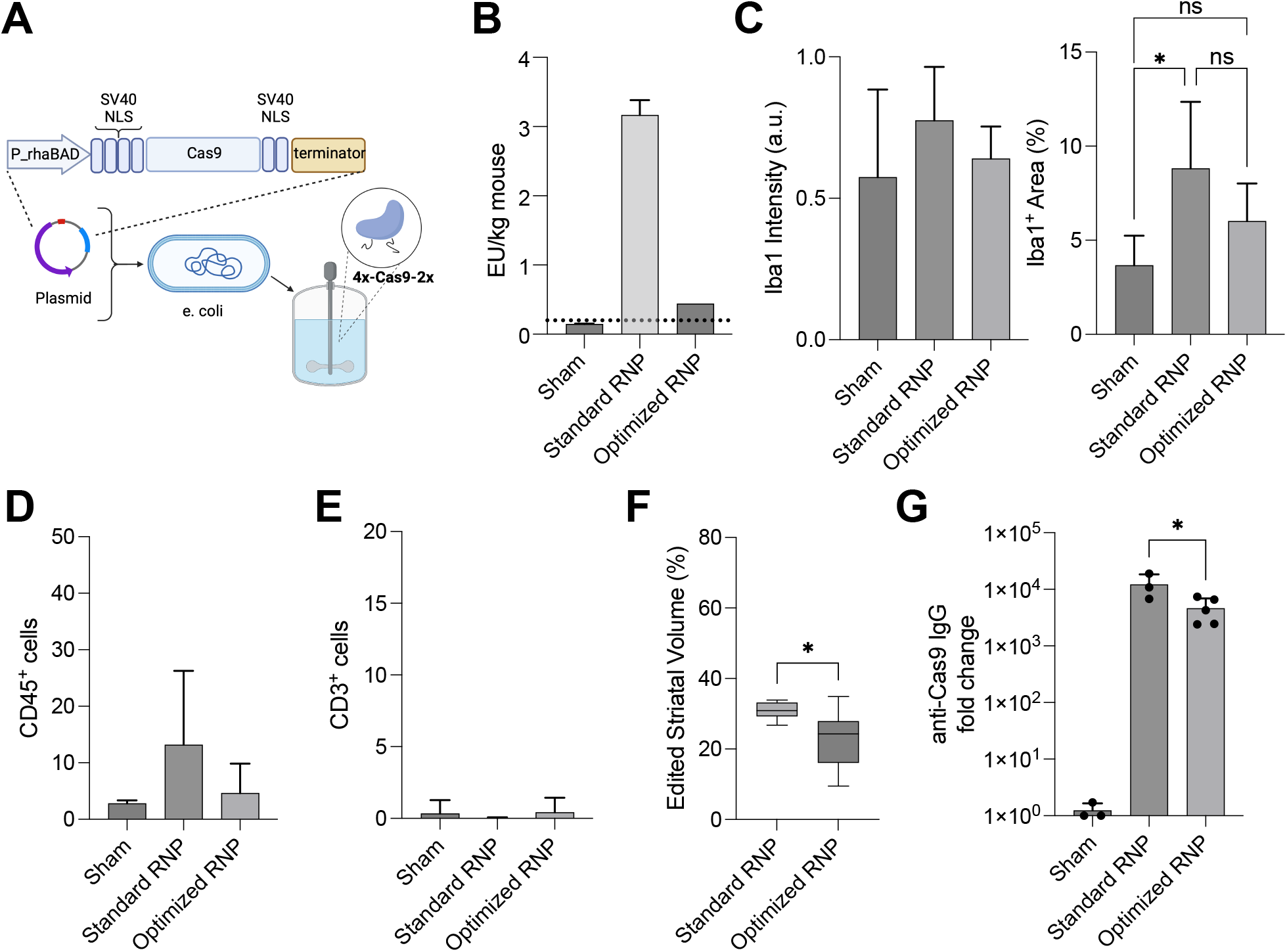
Optimized, low-endotoxin RNP formulation reduces local immune response. (A) Schematic of manufacturing scale up to produce industrial ultra-low endotoxin 4x-SpyCas9-2x protein using a tag-free expression and purification system. (B) Endotoxin levels calculated on a per mouse basis between the standard (laboratory 4x-SpyCas9-2x with sg298 2018) and optimized (industrial 4x-SpyCas9-2x protein with sg298 2022) RNP formulations at 25µM measured by LAL assay. The optimized RNP had a final endotoxin level of 0.44 EU/kg. Dotted line: FDA recommendation of 0.2 EU/kg/hr for drug products administered intrathecally in humans. (C) Quantification of Iba1^+^ staining intensity and percent area (n=6-10, one-way ANOVA, *p<0.05). (D) Quantification of CD45^+^ and (E) CD3^+^ cells per image (n=6-10, one-way ANOVA, ns). (F) Percent volume of edited striatal tissue for Cas9 RNPs injected at 25µM (n=6-10 injections). (G) Quantification of IgG antibodies against Cas9 or AAV capsid proteins measured 21-days after bilateral intrastriatal injections by ELISA (n=3-5 biological replicates).

Using the limulus amebocyte lysate (LAL) assay, we measured an endotoxin concentration of 0.035 EU/mg in the industrial-produced protein compared to 0.2 EU/mg in the lab-produced protein (Figure S12A). Interestingly, using the same assay, we found that guide RNA could be an unexpected source of endotoxin contamination. Endotoxin was present in at least three unopened vials of lyophilized RNA that had been stored at -80°C from a 2018 lot, but not in a more recently purchased lot from the same vendor when resuspended simultaneously (Figure S12A-C). To rule out false positives due to reaction of LAL with beta-glucans, we performed the Recombinant Factor C (rFC) assay. We measured a similar level of endotoxins in the guides between the LAL and rFC assays, demonstrating the positive signal was from contamination with endotoxin and not beta-glucans (Figure S12D-E).

To measure the physiological impact of endotoxin in our samples, we used HEK293 cells that were engineered to produce secreted embryonic alkaline phosphatase (SEAP) downstream of NF-κB activation resulting from human toll-like receptor 4 stimulation (hTLR4) with endotoxin/lipopolysaccharide (LPS, Figure S13A-E). The lab-produced protein stimulated NF-κB in HEK293 cells significantly greater than the industrial produced protein (p<0.01, unpaired t-test). Treatment with the industrially produced protein led to similar levels of SEAP between hTLR4 cells and the parental cell line (Null2), demonstrating that most of the NF-κB stimulation was downstream of other pattern-recognition receptors (such as TLR3, TLR5, or nucleotide-binding oligomerization domain-containing protein 1 (NOD-1) activation) and not due to LPS signaling through hTLR4 (Figure S13A-E). When combined with sg298 from the 2018 or 2022 lot, absorbance levels of SEAP further increased in RNP complexes made with lab produced protein, while the industrial protein with either guide did not induce a response (Figure S13F). Furthermore, guide RNA alone did not stimulate NF-κB in HEK293 cells (Figure S13D).

Finally, we measured endotoxins in the “optimized” formulation of RNPs, comprised of the industrially produced 4x-SpyCas9-2x protein and 2022 sgRNA, using the LAL assay. Estimating delivery of 10µL per mouse, the endotoxin burden was 0.44 EU/kg when RNPs were formulated at 25µM. These data suggest that RNPs could be delivered below the 0.2 EU/kg FDA threshold for intrathecal delivery^41^ when formulated at 10µM without significant loss of editing (Figure 3B and Figure 1H).

### Optimized RNP formulation reduces immune response

We performed CED bilateral intrastriatal injections to test if reducing endotoxins would improve the host immune response to RNPs *in vivo*. In this experiment, we compared the optimized RNP formulation (industrially produced 4x-SpyCas9-2x NLS protein with sg298 2022) to the standard formulation used in Figures 1 and 2 (laboratory produced 4x-SpyCas9-2x NLS protein with sg298 2018) at 25µM. The standard RNP induced a significant increase in Iba1^+^ area, consistent with our previous measurements (Figure 3C and Figure 1A); however, the optimized Cas9-RNP formulation did not induce microglial activation. Additionally, there was no increase in CD45^+^ and CD3^+^ cells from the sham in the optimized RNP group (Figure 3D). Of note, the standard RNP edited an average of 31 ± 3% striatal volume (greater than values reported in Figure 1, possibly due to differences between protein lots or injections), while the optimized RNP edited an average of 23 ± 8% striatal volume (Figure 3F). When tested *in vitro,* the optimized RNP performed slightly better at direct delivery than nucleofection compared to the standard formulation, which could explain in part the differences *in vivo* (Figure S14A). Interestingly, the standard RNP also lead to significantly greater anti-Cas9 IgG responses at three-weeks post-injection, possibly due to endotoxin boosting the adaptive immune response (Figure 3G). Taken together, we found that reducing endotoxins in both the guide RNA and protein components of the RNP leads to a reduced innate immune response, comparable to the sham, while maintaining high on-target editing. Furthermore, cell penetrant Cas9 proteins are amenable to expedited manufacturing of large quantities suitable for *in vivo* experiments.

In conclusion, our results establish complementary genome editing and immunogenicity outcomes between the two tested Cas9 delivery strategies (Figure S14B). To enable high-levels of editing in neurons within a localized brain region, minimizing adaptive immune responses, and timely and affordable manufacturing scale up, the RNP offers an effective alternative delivery system to viral vectors.

## Discussion

In the present study, we demonstrate that cell penetrant Cas9 RNPs edit a significant volume of the mouse striatum using convection enhanced delivery. Furthermore, the 4x-NLS modification enables self-delivery of Cas9 orthologous proteins *in vitro* to both mouse and human cells, demonstrating cross-species compatibility of the system for the first time. We also show that Cas9 RNPs have dose-dependent effects on the immune response, which can be mitigated by using ultra-low endotoxin protein produced in an industrial non-GMP setting. These experiments are informative for the design of future therapeutic applications of Cas9 RNP editors in mice and larger animal models.

Several studies have reported non-viral delivery of Cas9 into the mouse brain. The “CRISPR-Gold” Cas9 nanoparticle delivery system induced 14% edited glial cells near the injection site, sufficient to reduce repetitive behaviors in a mouse model of fragile X syndrome^42^. Additionally, incubating RNPs with R7L10, an arginine and leucine rich cationic peptide, induced 45% indels in the CA3 region of the hippocampus, leading to behavioral improvements in an Alzheimer’s disease mouse model^43^. Efficient editing of DARPP-32 medium spiny neurons in the striatum was achieved here and in recent work by others using RNPs packaged in biodegradable PEGylated nanocapsules^44^. Interestingly the PEGylated nanocapsules have a neutral charge, while 4x-Cas9-2x NLS RNPs have a net-positive charge, suggesting the mechanism of entry may differ between the two strategies. Systemic delivery of genome editors with glucose-conjugated silica nanoparticles and AAV9 can also lead to modest levels of editing in the brain, sufficient for therapeutic benefit ^45,46^. Despite the need for direct injection, the simplicity of the 4x-Cas9-2x RNP makes it ideal from a manufacturing perspective compared to other viral and nanoparticle formulations. The RNP formulation buffer could be further supplemented with polymers, such as polyethylene glycol (PEG)^47,48^, to possibly improve biodistribution with CED in the future.

Several studies show correlation between editing at the tdTomato locus and subsequent editing at endogenous sites^42,49,45^. It is important to note that expression of the tdTomato protein in the Ai9 mouse model underreports the actual genome editing efficiency, as double strand breaks that result in indels and small deletions are not sufficient to turn on the reporter^19^. We sought to further resolve editing outcomes with a ddPCR assay, since the tdTomato locus is too large for Illumina-based NGS. In the ddPCR probe drop-off assay we detected approximately a 6-fold increase in editing with AAV compared to RNP (approximately 15% versus 2.5% indels), whereas our immunofluorescent measurement showed a 2-fold increase. We speculate that the difference in reported editing efficiency between the two assays is based on the sampling methodology. The image analysis workflow quantified the volume of striatal tissue containing edited cells (Figure 1I), where 20-36% of neurons were edited within 25-50% of the striatum (Figure 1J). Therefore, the image quantification reflects the maximal biodistribution of edited cells in the striatum. The ddPCR assay utilized genomic DNA from 2-mm thick tissues dissected from the expected injection site including the striatum, as well as the cortex and corpus callosum, which likely favors editing events in the Cas9-AAV group due to its enhanced diffusion through the brain. While dose escalating the RNP did not improve editing levels, suggesting a saturating dose was already achieved as measured by image quantification, editing at different doses was not assessed using ddPCR. Furthermore, increasing the dose of Cas9-AAV may have further increased editing levels. Studies suggest that correcting pathological mutations in 20-30% of striatal neurons expressing mutant huntingtin protein is sufficient to significantly improve the disease pathology, therefore even modest editing levels in the striatum could enable therapeutic benefit^50^. Ultimately, behavioral assays are needed to further determine therapeutic benefit of the RNP approach in a disease-relevant model.

We hypothesized that cell penetrant Cas9-RNPs would be less immunogenic than Cas9-AAVs due to their transient expression. As the dose of 4x-SpyCas9-2x RNPs increased from 25µM to 100µM, there was an increase in CD45^+^ and GFAP^+^ cells, and a decrease in NeuN^+^ cells. As such, subsequent experiments were performed at 25µM, which was well-tolerated and resulted in similar levels of editing as the higher dose. The 25µM Cas9-RNP led to lower levels of vehicle-specific antibodies by 90-days post-injection compared to AAVs and did not upregulate gene signatures of T-cells at 28-days as measured by RT-qPCR, supporting our hypothesis. By 90 days, the levels of anti-Cas9 antibodies in the Cas9-AAV group were only elevated in three out of five mice, demonstrating that the kinetics of Cas9 antibody persistence were similar between RNPs and AAVs, despite stable, intracellular Cas9 expression. Reducing endotoxin in both the Cas9 protein and guide RNA prevented microglial reactions and reduced humoral responses at 21-days.

In the Cas9-AAV group, few immune cells (CD45, Iba1, or CD3) were observed in the striatum by immunostaining, however *CD3e* gene expression was significantly upregulated in explanted tissue, closely followed by an increase in *CD8a*. This finding could indicate accumulation of cytotoxic T-cells trafficking into the parenchyma from the blood vessels or ventricles. Additionally, no changes in NeuN, GFAP, and CD45 expression were observed in the Cas9-AAV group out to 4 months, demonstrating that the AAV delivery strategy was well-tolerated overall in naïve mice. A study by Li et al. found that mice immunized against SauCas9 with Freund’s adjuvant one week prior to intravenous delivery of AAV8-SauCas9-sgRNA resulted in accumulation of cytotoxic T-cells in the liver and subsequent removal of edited hepatocytes^12^. Therefore, the host immune response to Cas9-AAV in mice with pre-existing immunity would likely be different than what we observed in naïve mice. In the Cas9-RNP group, we found that pre-exposing mice to SpyCas9 protein with AddaVax^TM^ adjuvant 4 weeks prior to stereotaxic surgery synergistically increased systemic adaptive immune responses. Further studies are needed to assess the immune response to Cas9-AAV and RNP in models with pre-existing immunity, but how well these immunized mouse models recapitulate pre-existing immunity in humans is not clear. Furthermore, breakdown of the BBB in the context of neurodegenerative disease or strong expression of the tdTomato fluorescent reporter could also impact the host immune response^51,52^.

In this study, we used a strong CMV-promoter to drive expression of SauCas9 from the AAV, which allowed us to assess all subsets of edited cells in the striatum, in comparison to the RNP, which is not inherently designed to be neuron-specific. Although the SauCas9 transgene was still expressed 4-months post-delivery, editing at predicted off-target sites was not detected. Further work to experimentally determine guide-specific off-target sites, such as Guide-Seq^53^ or Circle-Seq^54^, was not performed. To prevent potential genotoxic side-effects due to long-term Cas9 expression, we recommend applying additional safeguards, such as AAV self-inactivation strategies and neuron-specific promoters, such as human synapsin 1 (hSyn), to increase cell specificity^55–57^. While self-inactivating AAVs may improve safety, they may not be sufficient to reduce partial integration of the viral genome at the Cas9 cut site, which has been reported^37–40^. Strategies to mitigate the host response to genome editors include providing immunosuppressants with CRISPR-Cas9 infusion and screening for pre-existing immunity prior to dosing when translating *in vivo* editing to humans^58^.

While SpyCas9 is generally the most efficacious and widely used Type II CRISPR protein to date, it is too large to be packaged in a single AAV with sgRNA. Therefore, in this study, we used AAV-CMV-SauCas9-U6-sgRNA as a positive control to benchmark delivery of RNPs, as similar AAV constructs have been used extensively in the literature and in clinical trials^24,23,12,59^. In vivo delivery of 4x-SauCas9-2x RNP did not lead to significant editing, thus, most comparisons were performed between two orthologous genome editors and delivery systems with different NLS configurations, which may impact the interpretation of this work. In vivo editing with 4x-SauCas9-2x RNP was possibly hindered by the reduced thermostability of the protein, as reported previously^60^. Employing Cas9 RNPs from a more thermostable organism could be advantageous in the future. Since RNPs are not restricted by cargo packaging, one advantage of the cell penetrant RNP technology is the ability to use SpyCas9 for future therapeutic applications in the brain.

In conclusion, the cell penetrant 4x-Cas9-2x NLS fusion protein enables simple and effective delivery of Cas9 RNPs into neurons *in vitro* and *in vivo* Our study is the first to comprehensively profile the host immune response to Cas9 in the brain, benchmark an RNP delivery strategy against the gold-standard for gene delivery in the CNS, and demonstrate feasibility of large-scale manufacturing. Given that Cas9-RNPs excel at editing high levels of neurons within a localized region of the brain, this is a promising modality to characterize therapeutic benefit in disease models in the future.

## Materials & Methods

### Plasmid construction

Cloning of several spacers into a plasmid encoding SauCas9 was performed as previously described. Oligonucleotides encoding sgRNAs were custom synthesized (Integrated DNA Technologies; IDT, Coralville, IA) and phosphorylated by T4 polynucleotide kinase (New England Biolabs; NEB, Ipswich, MA) for 30 min at 37°C. Oligonucleotides were annealed for 5 min at 95°C, cooled to room temperature and ligated into the BsmBI restriction sites of pSTX8,pKLT7.1_SaCas9prot_SaCas9guide plasmid. The following 23nt spacer sequences were cloned into the plasmid (spo 1: TGGTATGGCTGATTATGATCCTC; spo2: TCCCCCTGAACCTGAAACATAAA; spo3: GATGAGTTTGGACAAACCACAAC; spo4: TCCAGACATGATAAGATACATTG; spo5: CTCATCAATGTATCTTATCATGT), and plasmids were used for editing in mouse neural precursor cells *in vitro*. The best performing SauCas9 spacer (spo4: TCCAGACATGATAAGATACATTG) was then cloned into an AAV2 backbone plasmid. pX601-AAV-CMV::NLS-SaCas9-NLS-3xHA-bGHpA;U6::BsaI-sgRNA was a gift from Feng Zhang (Addgene plasmid # 61591; http://n2t.net/addgene:61591; RRID:Addgene_61591). Briefly, the plasmid was digested using BbsI and a pair of annealed oligos were cloned into the guide RNA destination site by Golden Gate assembly. Correct construction of all plasmids was verified by Sanger sequencing (UC Berkeley DNA Sequencing Facility).

### Recombinant adeno-associated virus (AAV) Production

The custom AAV9-CMV-61591-HA-Bgh vectors were produced at Virovek (Hayward, CA) in insect Sf9 cells by dual infection with rBV-inCap9-inRep-hr2 and rBV-CMV-61591-HA-Bgh. The AAV9-CMV-GFP vectors were produced by dual infection with rBVinCap9-inRep-hr2 and rBV-CMV-GFP. The vectors were purified through two rounds of cesium chloride (CsCl) ultracentrifugation. The CsCl was removed through buffer exchange with two PD-10 desalting columns. Viral titer (approximately 2e13 vg/mL) and purity were confirmed by nanodrop spectrophotometer, real-time PCR, and SDS-PAGE protein gel analysis. The vectors were passed through 0.2um sterilized filters, tested for endotoxins (< 0.6 EU/mL), as well as baculovirus and Sf9 DNA contamination (not detected).

### Purification of low-endotoxin proteins in a laboratory setting

Protein expression and purification was performed in the QB3 Macrolab at UC Berkeley using a custom low-endotoxin workflow. Briefly, the plasmid, 4xNLS-pMJ915v2 (Addgene plasmid # 88917; http://n2t.net/addgene:88917; RRID:Addgene_88917), was transformed into *E. coli* Rosetta2(DE3)pLysS cells (Novagen) and an overnight culture was used to inoculate 1 L flasks (12-24 L total per batch). Cells were grown for approximately 3 hours at 37°C then cooled to 16°C. At OD 0.8-0.9, cells were induced and harvested after 16-18 hours growth. Cells were lysed by homogenization in a buffer containing 1mM MgCl_2_ and benzonase (1:1000) to help reduce viscosity and centrifuged to remove insoluble material. Purification by Ni affinity (10 mL Ni resin for every 6 L cell lysate) was performed, and the bound protein was washed with 10 column volumes of buffer containing 0.1% Triton-X114 at 4°C to help reduce endotoxins. Tag removal with TEV protease (1:100) was performed overnight at 4°C, then heparin affinity was used to concentrate each batch of protein which was then flash frozen and stored at -80°C. A Sephacryl S300 size-exclusion column (SEC) and flow path were sanitized with 0.5 M NaOH overnight, then washed with up to 3 column volumes of buffer to rinse and equilibrate the system. Frozen samples were thawed, combined, and adjusted to 4.5 mL, and the S300 standard protocol was performed for size-exclusion. Samples were refrigerated overnight, and sanitation and size-exclusion were repeated the next day to further reduce endotoxin contamination. Peak fractions were pooled, concentrated to 40μM, aliquoted at 50μL, flash frozen in liquid nitrogen, and stored at -80°C in sterile, endotoxin-free Buffer 1 (25 mM NaP (pH 7.25), 300 mM NaCl, 200 mM trehalose (Sigma Aldrich #T5251, St. Louis, MO)). Final average protein yield was 1 mg per 1 L cells. Plasmids for 2xNLS-SauCas9-2xNLS, 3xNLS-SauCas9-2xNLS, and 4xNLS-SauCas9-2xNLS were created by deletion mutagenesis using the existing 4xNLS construct as a template. The genes were fully sequenced to confirm no additional mutations were introduced during the mutagenesis procedure.

### Purification of ultra-low endotoxin proteins in an industrial setting

4x-SpyCas9-2x NLS protein was manufactured according to Aldevron proprietary workflows for expression and purification of gene editing nucleases. Briefly, the gene for 4x-SpyCas9-2x NLS was synthesized (ATUM Bio, Sunnyvale, CA) and cloned into a pD881 expression vector (ATUM). Expression-ready plasmid DNA was transformed into E. coli BL21(DE3) (New England Biolabs) culture in animal-free TB media. At the appropriate OD600, expression was induced with 2.0% (w/v) Rhamnose and growth culture was harvested by centrifugation. Cells were lysed via dual-pass high-pressure homogenization and clarified via centrifugation. The clarified lysate was purified via multi-step chromatography using standard/commercially available resins. In the final chromatography step, the product is eluted via step elution and pooled to maximize final protein purity and minimize endotoxin. Product was dialyzed into the final formulation buffer, underwent three (3) exchanges of buffer, and was pooled into a sterile vessel for final filtration and dispensing. Product was evaluated for key quality attributes including endotoxin via PTS Endosafe assay (Charles River Labs, Cambridge, MA).

### Quantification of endotoxins in Cas9 RNPs

Proteins, guide RNAs, and pre-formed RNP complexes were subjected to several assays to quantify endotoxin burden according to the manufacturer’s instructions. All assays were performed with autoclaved or certified pyrogen-free plasticware and endotoxin (ET)-free water. The plate-reader based LAL assay was performed with the Endosafe Endochrome-K kit (Charles River, #R1708K), where a control standard endotoxin (CSE) was diluted from 5 EU/mL to 0.005 EU/mL. Samples were diluted 1:100 and plated in triplicate. An equal volume of LAL was added to each well. A Tecan Spark plate reader (Tecan, #30086376, M□nnendorf, Switzerland) with SparkControl magellan V 2.2 software was used at 37°C to read absorbance at 405nm every 30 seconds for 100 cycles. Time at which absorbance crossed optical density (OD) of 0.1 was recorded and used to determine endotoxin levels.

The cartridge-based LAL assay was performed using an Endosafe nexgen-PTS machine with R&D cartridges as recommended (Charles River, cat # PTS2005, 0.05 EU/mL sensitivity). Briefly, samples were diluted 1:50 in a large volume of ET-free water, vortexed, and 25μL was loaded into each of the four lanes of the cartridge, where two lanes contain CSE spike-in to calculate efficiency of the assay, which is valid from 50%-200% recovery. The final valid ET value was recorded from the duplicate measurement from a single cartridge.

The PyroGene Recombinant Factor C Endpoint Fluorescent Assay (Lonza, Walkersville, MD, cat # 50-658U) was performed as recommended. Kit-supplied CSE was diluted from 5 EU/mL to 0.005 EU/mL and samples were diluted 1:100 in ET-free water and added to a plate in triplicate. The plate was heated at 37°C for 10 minutes, then an equal volume of working reagent was added to each well. Fluorescence was read immediately at time 0 and again after incubating for 60 minutes. Relative fluorescence units (RFUs) of the ET-free water only blank wells were subtracted from all measurements, then delta RFUs between the two time points was calculated, and a linear regression was applied to the standard curve to calculate EUs in the samples. Fluorescence measurements were performed on a Cytation5 with Gen 5 3.04 software (BioTek, Winooski, VT).

HEK-Blue cells (hTLR4 and Null2) were purchased from Invivogen (San Diego, CA) and were grown under BSL2 conditions (37°C with 5% CO2) to measure SEAP production downstream of NFkappaB activation following treatment with Cas9 proteins, guide RNAs, and RNPs in vitro as recommended. Cells were grown in T-75 flasks with supplied antibiotic selection markers and passaged at 70% confluency. Cells were detached with gentle scraping in 1x PBS, centrifuged, counted, and plated for experiments in freshly prepared HEK-Blue Detection Media at approximately 32,000 cells per well in a 96-well plate. 180μL of cell suspension was plated directly into 20μL of diluted CSE (5 to 0.078 ng) or samples (diluted to 10μM) and incubated overnight at 37°C. Absorbance was read at 620nm in a Tecan Spark plate reader (Tecan, #30086376, MLnnendorf, Switzerland).

### Neural progenitor cell (NPC) line creation and culture

Neural progenitor cells were isolated from Ai9-tdTomato homozygous mouse embryos (day 13.5) by microdissection of cortical tissues into Hibernate E (#HECA, Brain Bits, LLC, Springfield, IL) and processing with the Neural Dissociation Kit with papain (#130-092-628, Miltenyi, Bergisch Gladbach, Germany) according to the manufacturer’s instruction. Single cells grew into non-adherent neurospheres, which were maintained in culture media (DMEM/F12 (ThermoFisher #10565-018, Waltham, MA), B-27 supplement (#12587-010), N-2 supplement (#17502-048), MEM non-essential amino acids (#11140-050), 10 mM HEPES (#15630-080), 1000X 2-mercaptoethanol (#21985-023), 100X Pen/Strep (#15140-122)) supplemented with growth factors (FGF-basic (Biolegend #579606) and EGF (ThermoFisher #PHG0311) to a final concentration of 20 ng/mL in media. Neurospheres were passaged every six days using the Neural Dissociation Kit to approximately 1.5 million cells per 10-cm dish and growth factors were refreshed every 3 days. Cells were authenticated by immunofluorescent staining for Nestin and GFAP, routinely tested for mycoplasma, and were used for experiments between passages 2 and 20. Dissociated cells were grown in monolayers in 96-well plates pre-coated with poly-DL-ornithine (SigmaAldrich, #P8638), laminin (SigmaAldrich #11243217001) and fibronectin (SigmaAldrich #F4759) at 10,000-30,000 cells per well for direct delivery and nucleofection experiments.

### Human induced pluripotent stem cell differentiation into NPCs and culture

MSC-iPSC1 cells were a generous gift from Boston Children’s Hospital. iPSCs were differentiated into NPCs based on dual SMAD inhibition as previously described. Briefly, iPSCs were plated onto Matrigel in the presence of 10μM Y-27632 (Sigma #Y0503) at a density of 200,000 cells/cm^2^. The next day (day 0) media was changed to KSR media (Knockout DMEM (ThermoFisher #10829018), 15% Knockout serum replacement (ThermoFisher #10828010), L-glutamine (1mM), 1% MEM Non-essential amino acids, and 0.1mM B-mercaptoethanol). Media was changed daily during differentiation and gradually changed from KSR media to N2/B27 media (Neurobasal medium (ThermoFisher #21103049), GlutaMAX Supplement (ThermoFisher #35050061), N-2 supplement (ThermoFisher #17502048) and B-27 supplement (ThermoFisher #17504044)) by increasing N2/B27 media to 1/3 on day 4, 2/3 on day 6 and full N2/B27 media on day 8. For the first 12 days of differentiation media was supplemented with 100nM LDN193189 (Sigma #SML0559) and 10μM SB431542 (Tocris Bioscience #1614, Bristol, England). For the first 4 days media was also supplemented with 2μM XAV939 (Tocris Bioscience #3748). On day 19, NPCs were dissociated with StemPro Accutase (ThermoFisher #A1110501) and replated onto Matrigel for expansion. NPCs were passaged every 6 days and maintained in NPC media (DMEM/F12, N2 supplement, B27 supplement and 20ng/ml bFGF (Corning, #354060, Corning, NY)) with media changes every other day. For direct delivery experiments, 12,000 cells were seeded in Matrigel in a 96-well plate and treated in triplicate with 100pmol of 4xNLS-SpyCas9-2xNLS RNPs with the EMX1 guide RNA (spacer: 5’ GAGTCCGAGCAGAAGAAGAA) or non-targeting guide RNA (spacer: 5’ AACGACTAGTTAGGCGTGTA). In the Lipofectamine^TM^ CRISPRmax group (ThermoFisher, #CMAX00003), 3μg of 0xNLS-SpyCas9-2xNLS protein (18 pmol) was mixed with sgRNA (1:1 molar ratio) in 8μL OptiMEM with 6μL of Cas9 Plus Reagent (1 μg protein: 2μL reagent) and was mixed with a second tube containing 3.6μL CRISPRmax reagent in 8μL OptiMEM, incubated for at least 5 minutes and was immediately distributed to cells in triplicate (1μg RNP per well), according to the manufacturer’s recommendations.

### Cas9 ribonucleoprotein (RNP) assembly and delivery to cells

For cell culture experiments, RNPs were prepared immediately before use at a 1.2:1 molar ratio of single guide RNA (Synthego, Redwood City, CA) to protein (QB3 Macrolab or Aldevron). The solution was incubated for 5-10 minutes at room temperature. For nucleofection, RNPs were formed at 10μM in 10μL of pre-supplemented buffer (Lonza P3 Primary Cell 96-well Kit, #V4SP-3096). A 15μL suspension of 250,000 mouse NPCs was mixed with 10μL RNP solution and added to the nucleofection cuvette. Nucleofection was performed using the 4D Nucleofector X Unit (Lonza, #AAF-1003X) with pulse code EH-100 and cells were recovered with 75 μL media per well approximately 2 minutes post-nucleofection. Nucleofected cells were then transferred to 100 μL fresh media in 96-well plates in triplicate and allowed to grow for 5 days at 37°C before analysis by flow cytometry for tdTomato expression. For direct delivery, RNPs were formed at 10μM in 10μL of sterile Buffer 1 (25 mM sodium phosphate pH (7.25), 100 mM NaCl, 200 mM trehalose). After NPCs were grown for two days in an adherent monolayer, 10μL was added to cell monolayer (“direct delivery”) for a final concentration of 1μM (100pmol RNP/100μL media). Media was changed 48-hours post-treatment and cells were collected 5 days post-treatment for analysis by flow cytometry for tdTomato expression or 4 days post-treatment for DNA sequencing.

For *in vivo* experiments, RNPs were prepared similarly at 10μM concentration in Buffer 1 and were incubated at 37°C for 10 minutes. RNPs were sterile filtered by centrifuging through 0.22μm Spin-X cellulose acetate membranes (Corning CoStar, #32119210) at 15,000xg for 1 minute at 4°C. RNPs were then concentrated using 100kDa Ultra-0.5 ml Centrifugal Filter Unit (Amicon, #, Burlington, MA) at 14,000xg at 4°C until the final desired concentration was reached (25-100μM, minimum 50μL volume) and collected by centrifuging at 1,000xg for 2 minutes. RNPs were then divided into single-use 20μL aliquots, flash frozen in liquid nitrogen, and stored at -80°C until the experiment. Prior to intracranial injection, RNPs were thawed, pipetted to mix, loaded into a 25μL syringe (Hamilton, #7654-01, Reno, NV) and injected with custom 29-gauge CED cannulas.

### AAV9 transduction

A single 50μL aliquot of AAV9-CMV-SauCas9-U6-sgRNA or AAV9-CMV-GFP (Virovek) was thawed from -80°C and stored at 4°C. AAV9 was diluted in 1x PBS without calcium or magnesium to the desired concentration. For *in vivo* experiments, AAV was diluted on the day of surgeries to 3e8-3e9 vg/μL and stored on ice until loaded into the syringe. 5μL was injected in each hemisphere to a final dose of 1.5e9-1.5e10 vg per hemisphere using the CED. For cell culture experiments, serial dilutions were performed from 1.6e9 vg/μL to 2e8 vg/μL (lowest MOI = 200,000) and 10μL of each were added into 96-well plates in triplicate and maintained for 3 days or 9 days prior to flow cytometry for GFP expression (transduction) and tdTomato expression (genome editing). AAV9 was added at the same time as NPC seeding for optimal transduction.

### Empty capsid quantification by cryo-electron microscopy

AAV samples were frozen using FEI Vitrobot Mark IV cooled down to 8°C at 100% humidity. Briefly, 4 µl of AAV9 capsids containing GFP or Cas9 cargo was deposited on 2/2 400 mesh C-flat grids (Electron Microscopy Sciences, Hatfield, PA, #CF224C-50), which were previously glow discharged at 15 mA for 15 s on PELCO easyGLOW instrument. Grids were blotted for 3 s with blot force 8 and wait time 2.5 s. Micrographs were collected manually on Talos Arctica operated at 200 kV and magnification 36,000x (pixel size 1.14 Å/pix) using a super-resolution camera setting (0.57 Å/pix) on K3 Direct Electron Detector. Micrographs were collected using SerialEM v. 3.8.7 software. Capids were counted manually by three blinded reviewers for each image and the three counts were averaged and reported as percentage empty capsids between the two groups.

### RNP size and aggregation measurement by dynamic light scattering (DLS)

RNPs were prepared at 25µM, according to the methods described above for *in vivo* experiments. Approximately 40µL of RNP was added to a disposable cuvette (ZEN0040) and inserted into the Zetasizer Nano (ZSP, Malvern, United Kingdom) then 173° scattering angle was measured at 25°C and again at 37°C after incubation for 30 minutes. Size distribution by mass histograms are shown with estimated peak size in nanometers (nm). Each sample was analyzed 3 times, resulting in a histogram generated from 10 individual measurements, for a total protocol run time of approximately 10 minutes.

### Analysis of editing in vitro

tdTomato positivity was assessed by flow cytometry using IGI facilities on the Attune NxT (Thermo Fisher, AFC2). Briefly, mouse NPCs were washed once with 1x PBS, harvested with 0.25% trypsin, neutralized with DMEM containing 10% FBS, and resuspended in 150uL of 1x PBS per well of a round-bottom 96-well plate for analysis. The percentage of tdTomato^+^ cells from each well was recorded. For analysis of genomic DNA (gDNA), media was removed, cells were rinsed once with 1X PBS, then incubated with 100 μL QuickExtract solution (Lucigen CorporationSupplier Diversity Partner QuickExtract DNA Extraction Solution 1.0, Fisher, #QE09050) at 37°C for 5 minutes. The cell lysate was then moved to a thermal cycler and incubated at 65°C for 20 minutes and 95°C for 20 minutes. gDNA was used in PCR reactions to generate amplicons of approximately 150-300bp for Illumina sequencing. A list of primers used for NGS is provided in Supplementary Table 1. Sequencing was performed with Illumina MiSeq in the IGI Center for Translational Genomics and reads were analyzed in CRISPResso (website).

### Stereotaxic infusion of Cas9 RNPs and AAVs

Ai9 mice (Jackson Laboratory, #007909, Bar Harbor, ME) were group housed at the University of California, Berkeley with a 12-hour light-dark cycle and allowed to feed and drink ad libitum. Housing, maintenance, and experimentation of the mice were carried out with strict adherence to ethical regulations set forth by the Animal Care and Use Committee (ACUC) at the University of California, Berkeley. Cas9-RNP and AAVs were prepared on-site at the University of California, Berkeley for injection into male and female tdTomato Ai9 mice between 2 to 5 months of age. All tools were autoclaved and injected materials were sterile. Mice anesthetized with 2% isoflurane, given pre-emptive analgesics, and were arranged on Angle-two stereotactic frame (Leica, Nussloch, Germany). The incision area was swabbed with three alternating wipes of 70% ethanol and betadine scrub with sterile applicators prior to performing minimally damaging craniotomies. The stereotaxic surgery coordinates used for targeting the striatum, relative to bregma, were +0.74 mm anteroposterior, ±1.90 mm mediolateral, −3.37 mm dorsoventral. Bilateral CED infusion of Cas9 RNPs (10-100μM) or Cas9 AAVs (3e8-3e9 vg/μL) was performed with a syringe pump to deliver 5μL at 0.5 μL per minute (Model 310 Plus, KD Scientific, Holliston, MA) with a step or non-step cannula. For intracerebroventricular (ICV) infusion of Cas9 RNPs, cannulas were placed at –0.7 mm anteroposterior, +/−1.2 mm mediolateral, and –2.5 mm dorsoventral according to Paxinos atlas of the adult mouse. Post-infusion, the syringes were left in position for 2 minutes before slow removal from the injection site, which was then cleaned, sutured, and surgically glued. Throughout the procedure, mice were kept at 37°C for warmth and Puralube Vet Ointment (Dechra, NDC #17033-211-38, Northwich, England) was applied to the outside of the eyes. For ICV injection of p0 neonatal mice, anesthesia was induced by hypothermia then 4μL of 100μM Cas9-RNP was injected with a hand-held 33-gauge needle unilaterally with 10% Fast Green dye to visualize distribution from one ventricle throughout the CNS. The needle was inserted 2 mm deep at a location approximately 0.25 mm lateral to the sagittal suture and 0.50-0.75 mm rostral to the neonatal coronal suture. RNP was slowly injected, then the needle was held in place for 15 seconds, and mice were monitored until recovery. For intrathecal injection, anesthetized mice received a 5, 25, or 50 μL bolus injection of Cas9 RNP at 300μM. The 29-gauge needle was inserted at the L6-S1 vertebral junction and angled slightly rostrally for the injection. Mice were allowed to fully recover before being transferred back to their housing. Recovery weight following all procedures was monitored daily for one week and mice were housed without further disruption for various time periods until tissue collection.

### Tissue collection and immunostaining

At the defined study endpoints (3, 21, and 90-days post-injection), mice were placed under anesthesia and tissues were perfused with 10mL of cold PBS and 5mL of 4% paraformaldehyde (PFA, Electron Microscopy Sciences, #15710, Hatfield, PA). Brains were post-fixed overnight in 4% PFA at 4°C, rinsed 3x with PBS, then cryoprotected in a 10% sucrose in PBS solution for approximately 3 days. Brains were embedded in optimal cutting temperature (OCT, Thermo Fisher, #23-730-571) media, and stored at -80°C. Brains were cut at 20-35 μm-thick sections using a cryostat (Leica CM3050S) and transferred to positively charged microscope slides. For immunohistochemical analysis, tissues were blocked with solution (0.3% TritonX-100, 1% bovine serum albumin (SigmaAldrich #A9418), 5% normal goat serum (SigmaAldrich, #G9023)) before 4°C incubation overnight with primary antibody in blocking solution. The next day, tissues were washed three times with PBS and incubated with secondary antibodies for one hour at room temperature. After three PBS washes, samples were incubated with DAPI solution (0.5 ug/mL, Roche LifeScience, Penzberg, Germany) as a DNA fluorescence probe for 10 minutes, washed three times with PBS, submerged once in deionized water, and mounted with glass coverslips in Fluoromount-G slide mounting medium (SouthernBiotech, Birmingham, AL). Primary antibodies included rabbit polyclonal anti-S100β (1:500, Abcam, #ab41548, Cambridge, England), rabbit polyclonal anti-Olig-2 (1:250, Millipore Sigma, #AB9610, Burlington, MA), rabbit polyclonal anti-doublecortin (1:800, Cell Signaling Technology, #4604, Danvers, MA), rabbit polyclonal anti-Ki67 (1:100, Abcam, #ab15580), mouse monoclonal anti-NeuN (1:500, Millipore Sigma, #MAB377), rabbit polyclonal anti-DARPP-32 (1:100, Cell Signaling Technology, #2302), rabbit polyclonal anti-Iba1 (1:100, Wako Chemicals, #019-19741, Richmond, VA), mouse monoclonal anti-glial fibrillary acidic protein (1:1000, Millipore Sigma, #MAB3402), rat monoclonal anti-CD45 (1:200, Thermo Fisher, #RA3-6B2), and rabbit polyclonal anti-CD3 (1:150, Abcam, #ab5690). Secondary antibodies included donkey anti-rat 488 (1:500, Thermo Fisher, #A-21208), goat anti-rabbit 488 (1:500, Thermo Fisher, #A32731), goat anti-rabbit 647 (1:500, Thermo Fisher, #A21245), and goat anti-mouse IgG1 647 (1:500, Thermo Fisher, #A-21240).

### Fluorescent imaging and image quantification

Whole brain sections were imaged and stitched using the automated AxioScanZ1 (Zeiss, Oberkochen, Germany) with a 20x objective in the DAPI and tdTomato channels. Images generated from slide scanning were viewed in ZenLite software (v 3.6 blue edition) as CZI files. Images were then exported to FIJI (v1.53q), Imaris (v 9.9.1), or QuPath (v 0.3.2) for quantification by authors blinded to the sample identity. The area of reflux from CED and blunt needles was calculated directly in ZenLite using the shape and area analysis tools. Immunostained cells and tissues were imaged on the Evos Revolve widefield microscope using a 20x objective or Stellaris 5 confocal microscope (Leica) with a 10x or 25x water-immersion objective to capture data in DAPI, FITC, tdTomato and CY5 channels. Approximately four images were taken at 20-25x per hemisphere across multiple sections for image quantification of CD45, Iba1, and CD3 (8-12 images quantified and averaged per injection). Approximately four to six z-stack images were captured and stitched per hemisphere for qualitative images of Iba1 and for quantification of NeuN, DARPP-32, ALDH1L1, and OLIG2 with tdTomato at 1024×1024 pixel resolution with a scanning speed of 100-200.

Measurements of striatal editing by volume were conducted using QuPath software (version 0.3.2) from images obtained from the Zeiss AxioscanZ1. Briefly, regions of interest (ROIs) were drawn to outline the border of each striatum and the inner area of tdTomato editing using the polygon tool to create annotations. All coronal plane areas were automatically calculated. Dorsoventral coordinates (relative to bregma) were then estimated in millimeters by consulting the Mouse Brain Atlas (C57BL/6J Coronal). Approximate tissue volume was calculated by averaging outlined areas between consecutive sections to represent the mean area across a dorsoventral segment and multiplying by the difference in dorsoventral coordinates. Edited striatal volumes were then divided by total striatal volumes to obtain percent editing. Additional tdTomato^+^ cell count measurements were conducted in Imaris software version 10.0 (Oxford Instruments, Abingdon, UK). Briefly, ROIs were drawn over each hemisphere (including cells in all brain sub-structures) using the “Segment only a Region of Interest” tool, and positive cells detected using the automated “Spots” tool to provide cell counts. Positivity thresholds were adjusted for each image to accurately capture edited cells manually. Counts of tdTomato cells on each image were then related back to approximate coordinates relative to bregma using the Mouse Brain Atlas (C57BL/6J Coronal) to quantify the distribution of edited cells.

Cell type specific measurements were conducted using QuPath software (version 0.3.2) on images obtained from Stellaris 5 z-stack maximal projections. ROIs were again drawn around areas of observed tdTomato editing, using the polygon tool to create a single annotation per image. Cell count calculations were performed using the “Cell Detection” and “Positive Cell Detection” tools, adjusting “Cell Mean” thresholds accordingly for each channel and image. Percent area and intensity measurements were performed in Fiji/ImageJ software (version 2.1.0/1.53c). Images were converted to 32-bit, and thresholds were adjusted to detect the corresponding stained area. Measurements were set to include area, minimum, maximum, and mean gray value, and area fraction, as well as to limit to threshold. All image quantification was performed on 2-5 serial sections with 3-10 independent injections per group for each analysis. Cell counts, area, intensity, and volume measurements were in general averaged from serial sections and were then grouped with other biological replicates, including independent injections, to report the treatment group average with standard deviation displayed by bar graph or box and whisker plot.

### Serum collection and ELISA

Blood was collected from mice at the time of euthanasia, allowed to clot at room temperature for 15-30 minutes, then centrifuged for 5 minutes at 2,000xg. Sera was collected and placed immediately on dry ice then stored at -80°C. Enzyme-linked immunosorbent assays were performed using the SeraCare Protein Detector™ HRP Microwell Anti-Mouse ELISA Kit, #5110-0011 (54-62-18) according to the manufacturer’s recommendations. First, 96-well plates were coated with antigens of interest (0.5 μg protein per well for SauCas9 and SpyCas9 (4xNLS protein variants) and approximately 1e9 empty AAV capsids per well) overnight at 4°C. Wells were washed three times and blocked at room temperature for one hour. Serum samples were then incubated in wells at varying concentrations (1:50 to 1:10,000 dilution) in 1X blocking buffer for four hours at room temperature, along with monoclonal antibody controls to generate a standard curve. Standards included CRISPR/Cas9 Monoclonal Antibody 7A9 (Epigentek, #A-9000-050, Farmingdale, NY), GenCRISPRLL SaCas9 Antibody 26H10 (GenScript, #A01952, Piscataway, NJ), and Anti-Adeno-associated Virus 9 Antibody clone HL2374 (Millipore Sigma, #MABF2326-25UG). Following three additional washes, the HRP secondary antibody was added at 1:500 in 1x blocking buffer and incubated for one hour. Wells were then washed three more times, and peroxidase substrate solutions were added. Absorbance was recorded at a wavelength of 405nm with Cytation5 plate reader with Gen 5 3.04 software (BioTek). Serum antibody concentrations were calculated using five-parameter logistic curve (5PL) data analysis at MyAssays.com and normalized to sham controls.

### Splenocyte collection and enzyme linked immunospot (ELISpot)

Spleens were collected at the time of euthanasia and stored in media composed of RPMI 1640 (ThermoFisher, #11875-119) with 10% FBS (VWR, #89510-186, Radnor, PA) and 1% P/S (ThermoFisher, #15140-122). Briefly, spleens were physically dissociated by forcing through a 100μm cell strainer in 10mL of media then single cells were passed through a 70μm strainer and centrifuged at 200xg for 5 minutes. Cells were resuspended in 5mL of 1x RBC Lysis Buffer (Miltenyi # 130-094-183) for approximately 3 minutes, then centrifuged again and resuspended in media for counting. The mouse interferon-gamma (IFN-Y) ELISpot kit (R&D Systems, #EL485, Minneapolis, MN) was used according to the manufacturer’s instructions to assess activation of splenocytes, containing T-cells, in response to treatment with Cas9 proteins. Briefly, the plate was pre-washed and 200μL of media for at least 20 minutes in the incubator, prior to adding cells at 300,000 per well in 100μL media. Treatments at 2x dose were prepared in media and 100μL was added to wells in triplicate (final 5 μg/mL concentration). Plates were wrapped in foil and incubated for 48 hours without disturbing. Concanavalin A (SigmaAldrich, #C5275) was used as a positive control for cell mediated IFN-Y production (final 4 μg/mL concentration). After 48-hours, cells were removed and the secreted analyte was detected with immunostaining using the kit-provided biotinylated monoclonal antibody specific for mouse IFN-Y, streptavidin-conjugated alkaline phosphatase, and stabilized detection mixture of 5-Bromo-4-Chloro-3’Indolylphosphate-p-Toluidine Salt (BCIP) and Nitro Blue Tetrazolium Chloride (NBT). After staining, plates were dried overnight and spot forming units were imaged and counted on the ImmunoSpot S6 Macro Analyzer (Cellular Technology Limited, Shaker Heights, OH).

### Immunization of mice to Cas9 with adjuvant

AddaVax™ (Invivogen, vac-adx-10), a squalene-based oil-in-water nano-emulsion, was mixed with an equal volume containing 25μg of 4x-SpyCas9-2x protein diluted in sterile buffer at room temperature for a final injection volume of 50μL. Mice received two 25μL subcutaneous injections of the AddaVax:Cas9 mixture (immunized) or AddaVax:Buffer alone (sham) with a 30-gauge insulin syringe into each flank. After four weeks, stereotaxic surgery with bilateral injections of 5μL of 25μM Cas9-RNPs was performed in a subset of mice. Mice showed no signs of pain or distress following treatment with AddaVax and no acute events were noted after surgery. Mice that received AddaVax, with or without surgery, were euthanized 6-weeks post-subcutaneous injections. Brains, serum, and spleens were collected for analysis of adaptive immune responses against repeated exposure to Cas9.

### DNA/RNA extraction from brain tissue slices and quantitative RT-PCR, droplet digital PCR, and long-read sequencing

Brains were collected at 3, 14, and 28-days or 4-months for DNA and RNA analysis. Briefly, mice were placed under anesthesia and perfused with cold PBS. Brains were harvested and cut into 2-mm sections using a matrix around the injection site (Zivic Instruments, Pittsburgh, PA). The slices were transferred onto chilled glass slides and further trimmed to approximate 30mg tissue weight (1–1.25 mm wide × 2 mm long). Tissues were flash frozen in liquid nitrogen then stored at -80°C until processing. DNA and RNA were collected from tissues using the AllPrep DNA/RNA Mini Kit (Qiagen, #80204, Venlo, Netherlands) according to the manufacturer’s instructions. Briefly, brains were homogenized in 1.5 mL tubes with a disposable pestle directly in RLT lysis buffer supplemented with 2-mercaptoethanol, then passed through Qiashredder columns to further homogenize prior to adding directly to the DNA and RNA binding columns. DNA was eluted in 100μL of EB, and RNA was eluted in 40 μL RNAse-free water. Concentrations of nucleic acids were measured by nanodrop spectrophotometer and samples were stored at -20°C.

Gene expression was quantified across multiple samples using a Custom RT^2^ PCR Array (Qiagen, #330171, CLAM45824) and analyzed using the RT² Profiler PCR Data Analysis Tool on GeneGlobe (Qiagen). For reverse transcription, the RT^2^ First Strand Kit (Qiagen, #330404) was used according to the manufacturer’s instructions. cDNA was diluted in water and added to the RT² SYBR Green qPCR Mastermix (Qiagen, #330502) then distributed across the 24-wells containing verified assay primers and controls (PCR array reproducibility control, reverse transcription efficiency control, genomic DNA contamination control, two house-keeping genes, and 19 experimental genes). Quantitative real-time PCR was performed on the CFX96 Touch Real-Time PCR System (BioRad). cDNA was also used in a droplet digital PCR reaction to measure SauCas9 expression at the 4-month time-point. qPCR assay IDs are included in Tables S1 and S2.

DNA was also used for PCR amplicon sequencing of predicted off-target sites and for droplet digital PCR (ddPCR). Off-target sites were predicted using Cas-OFFinder (http://www.rgenome.net/cas-offinder/)^61^. Predicted off-targets are described further in Tables S3 and S4. Primers were designed using NCBI Primer Blast with an amplicon size of 250-300bp, listed in Table S1. Sequencing was performed with Illumina MiSeq in the IGI Center for Translational Genomics and reads were analyzed in CRISPResso2 (http://crispresso.pinellolab.org)^62^.

For droplet digital PCR (ddPCR), a custom NHEJ ddPCR assays were generated using the online Bio-Rad design tool (Table S2). Assays for SauCas9 and SpyCas9 contain both the primers and probes (HEX-probe spanning the cut-site and a distal reference FAM-probe). To prepare the reactions, 110 ng of gDNA was combined with the 20x assay, 2x ddPCR Supermix for Probes (No dUTP), 1μL of of SmaI restriction enzyme (2 units per reaction), and water up to 22μL. Then 20 μl of each reaction mix was added to DG8TM Cartridges (Bio-Rad #1864008, Hercules, CA) followed by 70 μl of Droplet Generation Oil for Probes (Bio-Rad #1863005) and droplets were formed in the QX200 Droplet Generator. Droplets were then transferred to a 96-well plate and thermal cycled according to the manufacturer’s recommendation with a 3-minute annealing/extension step. After thermal cycling, the sealed plate was placed in the QX200 Droplet Reader and data was acquired and analyzed in the QuantaSoft Analysis Pro Software using the “Drop-Off” analysis, manually setting the thresholds for cluster calling (FAM+ only, FAM+ HEX+ cluster, FAM-HEX-cluster), and exporting fractional abundance calculations.

Long-read sequencing of the tdTomato locus was performed on DNA isolated from the treated mouse brains. Briefly, PCR amplicons were generated on 16 samples from the 4-month treatment groups using primers with unique barcodes for sample de-multiplexing. The KAPA HiFi Hotstart PCR Kit (Roche, KK2502) was used to amplify the 1100 bp product and reactions were cleaned with AMPure XP magnetic beads (Beckman Coulter Inc., Brea, CA) prior to analysis by Qubit and Bioanalyzer with the DNA 7500 Kit (Agilent, #5067-1506, Santa Clara, CA). Samples were combined and 1μg of pooled amplicons and submitted (> 20ng/μL) for sequencing with one PacBio Sequel 8M SMRT Cell at the QB3 Vincent Coates Genomic Sequencing Lab, yielding approximately 110,000 reads per sample. Data were analyzed using a custom pipeline to identify viral fragment trapping during DNA repair. Briefly, PacBio circular consensus reads were trimmed with Cutadapt (Version 4.1)^63^, then aligned to the AAV vector using NGMLR (Version 0.2.7)^64^ to generate BAM files. Soft-clipped regions of aligned reads were extracted using PySam (Version 0.18.0, https://github.com/pysam-developers/pysam) to parse CIGAR strings, then realigned to the tdTomato locus with NGMLR to verify integration within 200 bp of the cut site. Confirmed integrations were visualized along the AAV genome using pyGenomeTracks (Version 3.3) and coverage statistics were summarized using PySam^65,66^.

### Statistical analyses

The data presented in bar graphs and box and whisker plots are averages across multiple technical and biological replicates and error bars represent the standard deviation. Sample sizes are indicated in the text and figure legends and generally refer to technical injection replicates (two technical replicates, i.e., bilateral injections, per one biological replicate). When comparing two groups with normal distribution, an unpaired student’s t-test was performed in Prism 9 (GraphPad Software version 9.4.1). When comparing multiple groups, a one-way ANOVA with Tukey’s multiple comparison test was performed in Prism 9 (GraphPad Software version 9.4.1). The RT-qPCR experiments used Student’s t-test of the experimental group compared to the sham control (Qiagen GeneGlobe RT^2^ Profiler PCR Data Analysis). p ≤ 0.05 was considered significant

## Data Availability Statement

Long-read sequencing (BAM files from PacBio circular consensus sequence, CCS) are available in Sequence Read Archive (SRA). Accession number: [#]. All additional data is available upon request.

## Acknowledgements

Thank you to Netravathi Krishnappa and Christopher Hann-Soden for library preparation, sequencing, and analysis at the IGI Center for Translational Genomics and the QB3 Vincent J. Coates Genomics Sequencing Lab. Thank you to members of the Doudna lab at IGI, particularly Drs. Matthew Kan, Jennifer Hamilton, Cole Urnes, and Talia Wenger, as well as Brett Staahl, Ross Wilson, and Fyodor Urnov for intellectual contributions and support. Thank you to Aldevron, LLC for collaboration on this effort, especially Samantha Foti, Allison Pappas, David Yoder, and Max Sellman. Thank you to Denise Schichnes and Steven Ruzin at the CNR Berkeley Imaging Facility at the University of California, Berkeley as well as Feather Ives, Anita Flynn, and Holly Aaron at the CRL Microscopy Imaging Core (RRID:SCR_017852) at the University of California, Berkeley. Thank you to Eva Harris and lab members for the use of the ELISpot plate reader. Thank you to Kathy Snow and Ethan Saville at Jackson Laboratories for discussion about QuPath image quantification software. Thank you to the Bankiewicz lab, in particular Drs. Victor Van Laar, Lluis Samaranch, for discussion about CED. Thank you to Drs. Greg Barton and David Raulet for intellectual discussions regarding endotoxins and immune response. Thank you to Biogen Inc., especially Robin Kleinmen and Anirvan Ghosh, for funding and intellectual contributions. Thank you to Ruth L. Kirschstein F32 NIGMS for funding (E.C.S).

## Author Contributions

E.C.S performed the experiments, analysis, and manuscript preparation and oversaw contributions from researchers (R.A., E. A., S.E.K., A.S., N.L., Y.K.) who assisted with tissue processing, cloning, ELISAs, immunohistochemistry, and image analysis. E.C.S., J.K.S., and M.H.K. designed *in vivo* experiments, performed stereotaxic surgeries, confocal microscopy, and quantitative PCR. M.T. performed long-read NGS sequencing analysis. E.C.S. performed isolation and culture of mouse primary cells, flow cytometry, and ELISpot assays. K.S. performed electron microscopy. V.S.R. and L.T.V. performed human stem cell culture and differentiation. C.J., A.W., T.M., A.K., and T.F. performed custom low-endotoxin protein expression and purification. D.F.S. and J.A.D. approved the experiments, provided intellectual contributions, and co-wrote the manuscript.

## References

1. Heidenreich, M., and Zhang, F. (2016). Applications of CRISPR–Cas systems in neuroscience. Nat Rev Neurosci 17, 36–44. 10.1038/nrn.2015.2.

2. The SCGE Consortium, Saha, K., Sontheimer, E.J., Brooks, P.J., Dwinell, M.R., Gersbach, C.A., Liu, D.R., Murray, S.A., Tsai, S.Q., Wilson, R.C., et al. (2021). The NIH Somatic Cell Genome Editing program. Nature 592, 195–204. 10.1038/s41586-021-03191-1.

3. Jinek, M., Chylinski, K., Fonfara, I., Hauer, M., Doudna, J.A., and Charpentier, E. (2012). A Programmable Dual-RNA–Guided DNA Endonuclease in Adaptive Bacterial Immunity. Science 337, 816–821. 10.1126/science.1225829.

4. Jinek, M., East, A., Cheng, A., Lin, S., Ma, E., and Doudna, J. (2013). RNA-programmed genome editing in human cells. eLife 2, e00471. 10.7554/eLife.00471.

5. Doudna, J.A. (2020). The promise and challenge of therapeutic genome editing. Nature 578, 229–236. 10.1038/s41586-020-1978-5.

6. Hadaczek, P., Forsayeth, J., Mirek, H., Munson, K., Bringas, J., Pivirotto, P., L McBride, J., Davidson, B.L., and Bankiewicz, K.S. (2009). Transduction of Nonhuman Primate Brain with Adeno-Associated Virus Serotype 1: Vector Trafficking and Immune Response. Human Gene Therapy 20, 225–237. 10.1089/hum.2008.151.

7. Ciesielska, A., Hadaczek, P., Mittermeyer, G., Zhou, S., Wright, J.F., Bankiewicz, K.S., and Forsayeth, J. (2013). Cerebral Infusion of AAV9 Vector-encoding Non-self Proteins Can Elicit Cell-mediated Immune Responses. Molecular Therapy 21, 158–166. 10.1038/mt.2012.167.

8. Samaranch, L., Sebastian, W.S., Kells, A.P., Salegio, E.A., Heller, G., Bringas, J.R., Pivirotto, P., DeArmond, S., Forsayeth, J., and Bankiewicz, K.S. (2014). AAV9-mediated Expression of a Non-self Protein in Nonhuman Primate Central Nervous System Triggers Widespread Neuroinflammation Driven by Antigen-presenting Cell Transduction. Molecular Therapy 22, 329–337. 10.1038/mt.2013.266.

9. Agustín-Pavón, C., Mielcarek, M., Garriga-Canut, M., and Isalan, M. (2016). Deimmunization for gene therapy: host matching of synthetic zinc finger constructs enables long-term mutant Huntingtin repression in mice. Mol Neurodegeneration 11, 64. 10.1186/s13024-016-0128-x.

10. Chew, W.L., Tabebordbar, M., Cheng, J.K.W., Mali, P., Wu, E.Y., Ng, A.H.M., Zhu, K., Wagers, A.J., and Church, G.M. (2016). A multifunctional AAV–CRISPR–Cas9 and its host response. Nat Methods 13, 868–874. 10.1038/nmeth.3993.

11. Moreno, A.M., Palmer, N., Alemán, F., Chen, G., Pla, A., Jiang, N., Chew, W.L., Law, M., and Mali, P. (2019). immune-orthogonal orthologues of AAV capsids and of Cas9 circumvent the immune response to the administration of gene therapy. Nat Biomed Eng 3, 806–816. 10.1038/s41551-019-0431-2.

12. Li, A., Tanner, M.R., Lee, C.M., Hurley, A.E., De Giorgi, M., Jarrett, K.E., Davis, T.H., Doerfler, A.M., Bao, G., Beeton, C., et al. (2020). AAV-CRISPR Gene Editing Is Negated by Pre-existing Immunity to Cas9. Molecular Therapy 28, 1432–1441. 10.1016/j.ymthe.2020.04.017.

13. Simhadri, V.L., McGill, J., McMahon, S., Wang, J., Jiang, H., and Sauna, Z.E. (2018). Prevalence of Pre-existing Antibodies to CRISPR-Associated Nuclease Cas9 in the USA Population. Molecular Therapy - Methods & Clinical Development 10, 105–112. 10.1016/j.omtm.2018.06.006.

14. Charlesworth, C.T., Deshpande, P.S., Dever, D.P., Camarena, J., Lemgart, V.T., Cromer, M.K., Vakulskas, C.A., Collingwood, M.A., Zhang, L., Bode, N.M., et al. (2019). Identification of preexisting adaptive immunity to Cas9 proteins in humans. Nat Med 25, 249–254. 10.1038/s41591-018-0326-x.

15. Wagner, D.L., Amini, L., Wendering, D.J., Burkhardt, L.-M., Akyüz, L., Reinke, P., Volk, H.-D., and Schmueck-Henneresse, M. (2019). High prevalence of Streptococcus pyogenes Cas9-reactive T cells within the adult human population. Nat Med 25, 242–248. 10.1038/s41591-018-0204-6.

16. Ferdosi, S.R., Ewaisha, R., Moghadam, F., Krishna, S., Park, J.G., Ebrahimkhani, M.R., Kiani, S., and Anderson, K.S. (2019). Multifunctional CRISPR-Cas9 with engineered immunosilenced human T cell epitopes. Nat Commun 10, 1842. 10.1038/s41467-019-09693-x.

17. Tang, X.-Z.E., Tan, S.X., Hoon, S., and Yeo, G.W. (2022). Pre-existing adaptive immunity to the RNA-editing enzyme Cas13d in humans. Nat Med 28, 1372–1376. 10.1038/s41591-022-01848-6.

18. Mingozzi, F., and High, K.A. (2013). Immune responses to AAV vectors: overcoming barriers to successful gene therapy. Blood 122, 23–36. 10.1182/blood-2013-01-306647.

19. Staahl, B.T., Benekareddy, M., Coulon-Bainier, C., Banfal, A.A., Floor, S.N., Sabo, J.K., Urnes, C., Munares, G.A., Ghosh, A., and Doudna, J.A. (2017). Efficient genome editing in the mouse brain by local delivery of engineered Cas9 ribonucleoprotein complexes. Nat Biotechnol 35, 431–434. 10.1038/nbt.3806.

20. Liu, J., Gaj, T., Wallen, M.C., and Barbas, C.F. (2015). Improved Cell-Penetrating Zinc-Finger Nuclease Proteins for Precision Genome Engineering. Molecular Therapy - Nucleic Acids 4, e232. 10.1038/mtna.2015.6.

21. Madisen, L., Zwingman, T.A., Sunkin, S.M., Oh, S.W., Zariwala, H.A., Gu, H., Ng, L.L., Palmiter, R.D., Hawrylycz, M.J., Jones, A.R., et al. (2010). A robust and high-throughput Cre reporting and characterization system for the whole mouse brain. Nat Neurosci 13, 133– 140. 10.1038/nn.2467.

22. Ran, F.A., Cong, L., Yan, W.X., Scott, D.A., Gootenberg, J.S., Kriz, A.J., Zetsche, B., Shalem, O., Wu, X., Makarova, K.S., et al. (2015). In vivo genome editing using Staphylococcus aureus Cas9. Nature 520, 186–191. 10.1038/nature14299.

23. Maeder, M.L., Stefanidakis, M., Wilson, C.J., Baral, R., Barrera, L.A., Bounoutas, G.S., Bumcrot, D., Chao, H., Ciulla, D.M., DaSilva, J.A., et al. (2019). Development of a gene-editing approach to restore vision loss in Leber congenital amaurosis type 10. Nat Med 25, 229–233. 10.1038/s41591-018-0327-9.

24. Yin, C., Zhang, T., Qu, X., Zhang, Y., Putatunda, R., Xiao, X., Li, F., Xiao, W., Zhao, H., Dai, S., et al. (2017). In Vivo Excision of HIV-1 Provirus by saCas9 and Multiplex Single-Guide RNAs in Animal Models. Molecular Therapy 25, 1168–1186. 10.1016/j.ymthe.2017.03.012.

25. Gao, G., Vandenberghe, L.H., Alvira, M.R., Lu, Y., Calcedo, R., Zhou, X., and Wilson, J.M. (2004). Clades of Adeno-Associated Viruses Are Widely Disseminated in Human Tissues. J Virol 78, 6381–6388. 10.1128/JVI.78.12.6381-6388.2004.

26. Smith, R.H., Levy, J.R., and Kotin, R.M. (2009). A Simplified Baculovirus-AAV Expression Vector System Coupled With One-step Affinity Purification Yields High-titer rAAV Stocks From Insect Cells. Molecular Therapy 17, 1888–1896. 10.1038/mt.2009.128.

27. Chen, C., Akerstrom, V., Baus, J., Lan, M.S., and Breslin, M.B. (2013). Comparative analysis of the transduction efficiency of five adeno associated virus serotypes and VSV-G pseudotype lentiviral vector in lung cancer cells. Virol J 10, 86. 10.1186/1743-422X-10-86.

28. Gaj, T., Staahl, B.T., Rodrigues, G.M.C., Limsirichai, P., Ekman, F.K., Doudna, J.A., and Schaffer, D.V. (2017). Targeted gene knock-in by homology-directed genome editing using Cas9 ribonucleoprotein and AAV donor delivery. Nucleic Acids Research 45, e98–e98. 10.1093/nar/gkx154.

29. Park, I.-H., Zhao, R., West, J.A., Yabuuchi, A., Huo, H., Ince, T.A., Lerou, P.H., Lensch, M.W., and Daley, G.Q. (2008). Reprogramming of human somatic cells to pluripotency with defined factors. Nature 451, 141–146. 10.1038/nature06534.

30. Qi, Y., Zhang, X.-J., Renier, N., Wu, Z., Atkin, T., Sun, Z., Ozair, M.Z., Tchieu, J., Zimmer, B., Fattahi, F., et al. (2017). Combined small-molecule inhibition accelerates the derivation of functional cortical neurons from human pluripotent stem cells. Nat Biotechnol 35, 154–163. 10.1038/nbt.3777.

31. Chambers, S.M., Fasano, C.A., Papapetrou, E.P., Tomishima, M., Sadelain, M., and Studer, L. (2009). Highly efficient neural conversion of human ES and iPS cells by dual inhibition of SMAD signaling. Nat Biotechnol 27, 275–280. 10.1038/nbt.1529.

32. Fu, Y., Foden, J.A., Khayter, C., Maeder, M.L., Reyon, D., Joung, J.K., and Sander, J.D. (2013). High-frequency off-target mutagenesis induced by CRISPR-Cas nucleases in human cells. Nat Biotechnol 31, 822–826. 10.1038/nbt.2623.

33. Bobo, R.H., Laske, D.W., Akbasak, A., Morrison, P.F., Dedrick, R.L., and Oldfield, E.H. (1994). Convection-enhanced delivery of macromolecules in the brain. Proc. Natl. Acad. Sci. U.S.A. 91, 2076–2080. 10.1073/pnas.91.6.2076.

34. Hadaczek, P., Kohutnicka, M., Krauze, M.T., Bringas, J., Pivirotto, P., Cunningham, J., and Bankiewicz, K. (2006). Convection-Enhanced Delivery of Adeno-Associated Virus Type 2 (AAV2) into the Striatum and Transport of AAV2 Within Monkey Brain. Human Gene Therapy 17, 291–302. 10.1089/hum.2006.17.291.

35. Krauze, M.T., Saito, R., Noble, C., Tamas, M., Bringas, J., Park, J.W., Berger, M.S., and Bankiewicz, K. (2005). Reflux-free cannula for convection-enhanced high-speed delivery of therapeutic agents: Technical note. Journal of Neurosurgery 103, 923–929. 10.3171/jns.2005.103.5.0923.

36. Aschauer, D.F., Kreuz, S., and Rumpel, S. (2013). Analysis of Transduction Efficiency, Tropism and Axonal Transport of AAV Serotypes 1, 2, 5, 6, 8 and 9 in the Mouse Brain. PLoS ONE 8, e76310. 10.1371/journal.pone.0076310.

37. Hanlon, K.S., Kleinstiver, B.P., Garcia, S.P., Zaborowski, M.P., Volak, A., Spirig, S.E., Muller, A., Sousa, A.A., Tsai, S.Q., Bengtsson, N.E., et al. (2019). High levels of AAV vector integration into CRISPR-induced DNA breaks. Nat Commun 10, 4439. 10.1038/s41467-019-12449-2.

38. Nelson, C.E., Wu, Y., Gemberling, M.P., Oliver, M.L., Waller, M.A., Bohning, J.D., Robinson-Hamm, J.N., Bulaklak, K., Castellanos Rivera, R.M., Collier, J.H., et al. (2019). Long-term evaluation of AAV-CRISPR genome editing for Duchenne muscular dystrophy. Nat Med 25, 427–432. 10.1038/s41591-019-0344-3.

39. Ferrari, S., Jacob, A., Cesana, D., Laugel, M., Beretta, S., Varesi, A., Unali, G., Conti, A., Canarutto, D., Albano, L., et al. (2022). Choice of template delivery mitigates the genotoxic risk and adverse impact of editing in human hematopoietic stem cells. Cell Stem Cell 29, 1428–1444.e9. 10.1016/j.stem.2022.09.001.

40. Simpson, B.P., Yrigollen, C.M., Izda, A., and Davidson, B.L. (2023). Targeted long-read sequencing captures CRISPR editing and AAV integration outcomes in brain. Molecular Therapy, S1525001623000047. 10.1016/j.ymthe.2023.01.004.

41. Zink McCullough, K. Calculating Endotoxin Limits for Drug Products (American Pharmaceutical Review).

42. Lee, B., Lee, K., Panda, S., Gonzales-Rojas, R., Chong, A., Bugay, V., Park, H.M., Brenner, R., Murthy, N., and Lee, H.Y. (2018). Nanoparticle delivery of CRISPR into the brain rescues a mouse model of fragile X syndrome from exaggerated repetitive behaviours. Nat Biomed Eng 2, 497–507. 10.1038/s41551-018-0252-8.

43. Park, H., Oh, J., Shim, G., Cho, B., Chang, Y., Kim, S., Baek, S., Kim, H., Shin, J., Choi, H., et al. (2019). In vivo neuronal gene editing via CRISPR–Cas9 amphiphilic nanocomplexes alleviates deficits in mouse models of Alzheimer’s disease. Nat Neurosci 22, 524–528. 10.1038/s41593-019-0352-0.

44. Metzger, J.M., Wang, Y., Neuman, S.S., Snow, K.J., Murray, S.A., Lutz, C.M., Bondarenko, V., Felton, J., Gimse, K., Xie, R., et al. (2023). Efficient in vivo neuronal genome editing in the mouse brain using nanocapsules containing CRISPR-Cas9 ribonucleoproteins. Biomaterials 293, 121959. 10.1016/j.biomaterials.2022.121959.

45. Wang, Y., Wang, X., Xie, R., Burger, J.C., Tong, Y., and Gong, S. (2022). Overcoming the Blood–Brain Barrier for Gene Therapy via Systemic Administration of GSH-Responsive Silica Nanocapsules. Advanced Materials, 2208018. 10.1002/adma.202208018.

46. Yan, S., Zheng, X., Lin, Y., Li, C., Liu, Z., Li, J., Tu, Z., Zhao, Y., Huang, C., Chen, Y., et al. (2023). Cas9-mediated replacement of expanded CAG repeats in a pig model of Huntington’s disease. Nat. Biomed. Eng. 10.1038/s41551-023-01007-3.

47. He, M., Li, J., Han, H., Borges, C.A., Neiman, G., Røise, J.J., Hadaczek, P., Mendonsa, R., Holm, V.R., Wilson, R.C., et al. (2020). A traceless linker for aliphatic amines that rapidly and quantitatively fragments after reduction. Chem. Sci. 11, 8973–8980. 10.1039/D0SC00929F.

48. Saito, R., Krauze, M.T., Noble, C.O., Tamas, M., Drummond, D.C., Kirpotin, D.B., Berger, M.S., Park, J.W., and Bankiewicz, K.S. (2006). Tissue affinity of the infusate affects the distribution volume during convection-enhanced delivery into rodent brains: Implications for local drug delivery. Journal of Neuroscience Methods 154, 225–232. 10.1016/j.jneumeth.2005.12.027.

49. Wei, T., Cheng, Q., Min, Y.-L., Olson, E.N., and Siegwart, D.J. (2020). Systemic nanoparticle delivery of CRISPR-Cas9 ribonucleoproteins for effective tissue specific genome editing. Nat Commun 11, 3232. 10.1038/s41467-020-17029-3.

50. Oura, S., Noda, T., Morimura, N., Hitoshi, S., Nishimasu, H., Nagai, Y., Nureki, O., and Ikawa, M. (2021). Precise CAG repeat contraction in a Huntington’s Disease mouse model is enabled by gene editing with SpCas9-NG. Commun Biol 4, 771. 10.1038/s42003-021-02304-w.

51. Sweeney, M.D., Sagare, A.P., and Zlokovic, B.V. (2018). Blood–brain barrier breakdown in Alzheimer disease and other neurodegenerative disorders. Nat Rev Neurol 14, 133–150. 10.1038/nrneurol.2017.188.

52. Huang, L., Bommireddy, R., Munoz, L.E., Guin, R.N., Wei, C., Ruggieri, A., Menon, A.P., Li, X., Shanmugam, M., Owonikoko, T.K., et al. (2021). Expression of tdTomato and luciferase in a murine lung cancer alters the growth and immune microenvironment of the tumor. PLoS ONE 16, e0254125. 10.1371/journal.pone.0254125.

53. Tsai, S.Q., Zheng, Z., Nguyen, N.T., Liebers, M., Topkar, V.V., Thapar, V., Wyvekens, N., Khayter, C., Iafrate, A.J., Le, L.P., et al. (2015). GUIDE-seq enables genome-wide profiling of off-target cleavage by CRISPR-Cas nucleases. Nat Biotechnol 33, 187–197. 10.1038/nbt.3117.

54. Tsai, S.Q., Nguyen, N.T., Malagon-Lopez, J., Topkar, V.V., Aryee, M.J., and Joung, J.K. (2017). CIRCLE-seq: a highly sensitive in vitro screen for genome-wide CRISPR–Cas9 nuclease off-targets. Nat Methods 14, 607–614. 10.1038/nmeth.4278.

55. Ibraheim, R., Tai, P.W.L., Mir, A., Javeed, N., Wang, J., Rodríguez, T.C., Namkung, S., Nelson, S., Khokhar, E.S., Mintzer, E., et al. (2021). Self-inactivating, all-in-one AAV vectors for precision Cas9 genome editing via homology-directed repair in vivo. Nat Commun 12, 6267. 10.1038/s41467-021-26518-y.

56. Li, A., Lee, C.M., Hurley, A.E., Jarrett, K.E., De Giorgi, M., Lu, W., Balderrama, K.S., Doerfler, A.M., Deshmukh, H., Ray, A., et al. (2019). A Self-Deleting AAV-CRISPR System for In Vivo Genome Editing. Mol Ther Methods Clin Dev 12, 111–122. 10.1016/j.omtm.2018.11.009.

57. Merienne, N., Vachey, G., de Longprez, L., Meunier, C., Zimmer, V., Perriard, G., Canales, M., Mathias, A., Herrgott, L., Beltraminelli, T., et al. (2017). The Self-Inactivating KamiCas9 System for the Editing of CNS Disease Genes. Cell Rep 20, 2980–2991. 10.1016/j.celrep.2017.08.075.

58. Ewaisha, R., and Anderson, K.S. (2023). Immunogenicity of CRISPR therapeutics—Critical considerations for clinical translation. Front. Bioeng. Biotechnol. 11, 1138596. 10.3389/fbioe.2023.1138596.

59. Mancuso, P., Chen, C., Kaminski, R., Gordon, J., Liao, S., Robinson, J.A., Smith, M.D., Liu, H., Sariyer, I.K., Sariyer, R., et al. (2020). CRISPR based editing of SIV proviral DNA in ART treated non-human primates. Nat Commun 11, 6065. 10.1038/s41467-020-19821-7.

60. Harrington, L.B., Paez-Espino, D., Staahl, B.T., Chen, J.S., Ma, E., Kyrpides, N.C., and Doudna, J.A. (2017). A thermostable Cas9 with increased lifetime in human plasma. Nat Commun 8, 1424. 10.1038/s41467-017-01408-4.

61. Bae, S., Park, J., and Kim, J.-S. (2014). Cas-OFFinder: a fast and versatile algorithm that searches for potential off-target sites of Cas9 RNA-guided endonucleases. Bioinformatics 30, 1473–1475. 10.1093/bioinformatics/btu048.

62. Clement, K., Rees, H., Canver, M.C., Gehrke, J.M., Farouni, R., Hsu, J.Y., Cole, M.A., Liu, D.R., Joung, J.K., Bauer, D.E., et al. (2019). CRISPResso2 provides accurate and rapid genome editing sequence analysis. Nat Biotechnol 37, 224–226. 10.1038/s41587-019-0032-3.

63. Martin, M. (2011). Cutadapt removes adapter sequences from high-throughput sequencing reads. EMBnet j. 17, 10. 10.14806/ej.17.1.200.

64. Sedlazeck, F.J., Rescheneder, P., Smolka, M., Fang, H., Nattestad, M., von Haeseler, A., and Schatz, M.C. (2018). Accurate detection of complex structural variations using single-molecule sequencing. Nat Methods 15, 461–468. 10.1038/s41592-018-0001-7.

65. Ramírez, F., Bhardwaj, V., Arrigoni, L., Lam, K.C., Grüning, B.A., Villaveces, J., Habermann, B., Akhtar, A., and Manke, T. (2018). High-resolution TADs reveal DNA sequences underlying genome organization in flies. Nat Commun 9, 189. 10.1038/s41467-017-02525-w.

66. Lopez-Delisle, L., Rabbani, L., Wolff, J., Bhardwaj, V., Backofen, R., Grüning, B., Ramírez, F., and Manke, T. (2021). pyGenomeTracks: reproducible plots for multivariate genomic datasets. Bioinformatics 37, 422–423. 10.1093/bioinformatics/btaa692.

