## Supplementary files for "Genome editing in the mouse brain with minimally immunogenic Cas9 RNPs"

**Supplemental Table 1.**

Primer sequences for NGS amplicon sequencing and ddPCR. All primers for Illumina MiSeq were ordered with (GCTCTTCCGATCT) at the 5’ end for library preparation and indexing.

| **Primer Name** | **Sequence (5’ → 3’)** | **Assay/Application** |
|---|---|---|
| BS-272 Forward | GCTCCTGGGCAACGTGCTGGTTATTG | tdTomato on-target (long read PacBio) |
| BS-273 Reverse | TTGATGACCTCCTCGCCCTTGCTCAC | tdTomato on-target (long read PacBio) |
| SauCas9 OT1 (chr2) Forward | GCACATGGACATGGATTTGTTCA | tdTomato off-target (short read Illumina MiSeq) |
| SauCas9 OT1 (chr2) Reverse | AGACCCACAAAGATCACAGGTAA | tdTomato off-target (short read Illumina MiSeq) |
| SauCas9 OT2 (chr18) Forward | TCATCTTTTGGGGGATTGCCT | tdTomato off-target (short read Illumina MiSeq) |
| SauCas9 OT2 (chr18) Reverse | AGGCCATTGTCCATGGAGTC | tdTomato off-target (short read Illumina MiSeq) |
| SauCas9 OT3 (chr1) Forward | CATGCTTACCACAGGCTCCA | tdTomato off-target (short read Illumina MiSeq) |
| SauCas9 OT3 (chr1) Reverse | ACTGTTACCCAGCCTCTCCT | tdTomato off-target (short read Illumina MiSeq) |
| SauCas9 OT4 (chr13) Forward | TTTAAGGACTTGGCAGACCACT | tdTomato off-target (short read Illumina MiSeq) |
| SauCas9 OT4 (chr13) Reverse | TGAGCGACCATGACCCTGTAA | tdTomato off-target (short read Illumina MiSeq) |
| SauCas9 OT5 (chr17) Forward | TGCCAACAGAAAAACCACAGC | tdTomato off-target (short read Illumina MiSeq) |
| SauCas9 OT5 (chr17) Reverse | AGCTTCCCTAAACCCAAGAGC | tdTomato off-target (short read Illumina MiSeq) |
| SauCas9 OT6 (chr4-1) Forward | TCACCCAGCAACTTGTGGAA | tdTomato off-target (short read Illumina MiSeq) |
| SauCas9 OT6 (chr4-1) Reverse | TAGGGATCCAAAAGCTGGGA | tdTomato off-target (short read Illumina MiSeq) |
| SauCas9 OT7 (chr4-2) Forward | GCAGAGCAGCAGGCATTCTT | tdTomato off-target (short read Illumina MiSeq) |
| SauCas9 OT7 (chr4-2) Reverse | AGGTTCACCCATTCTTGACTTCT | tdTomato off-target (short read Illumina MiSeq) |
| SauCas9 OT8 (chr9) Forward | TCAGAGACAACAATCCTAGCAGA | tdTomato off-target (short read Illumina MiSeq) |
| SauCas9 OT8 (chr9) Reverse | GGGCCTATCTGTCCTTGGGTA | tdTomato off-target (short read Illumina MiSeq) |
| SauCas9 OT9 (chrX-1) Forward | GGTTCCAAACCTCCCTAACAAC | tdTomato off-target (short read Illumina MiSeq) |
| SauCas9 OT9 (chrX-1) Reverse | ATGTCATCCCTAGTTCCTATGTAAA | tdTomato off-target (short read Illumina MiSeq) |
| SauCas9 OT10 (chrX-2) Forward | CCCTGGGCTTAGCCATTTCT | tdTomato off-target (short read Illumina MiSeq) |

| | | |
|---|---|---|
| SauCas9 OT10 (chrX-2) Reverse | CCCACAACCCTAGTTGAGCC | tdTomato off-target (short read Illumina MiSeq) |
| SpyCas9 OT1 (chr4-3) Forward | CTGGTTCCCACTGGACCTTC | tdTomato off-target (short read Illumina MiSeq) |
| SpyCas9 OT1 (chr4-3) Reverse | GCCATGGTGTGTAAAGTAGGTG | tdTomato off-target (short read Illumina MiSeq) |
| SpyCas9 OT2 (chr17-2) Forward | ATCCATGTTGGGGGTTTAGTT | tdTomato off-target (short read Illumina MiSeq) |
| SpyCas9 OT2 (chr17-2) Reverse | GGGGGACTTTTGGGATAGCA | tdTomato off-target (short read Illumina MiSeq) |
| SpyCas9 OT3 (chr12) Forward | CCACATTCCCATCCCCAGAA | tdTomato off-target (short read Illumina MiSeq) |
| SpyCas9 OT3 (chr12) Reverse | CAGTTTTGCTAGGGGGAGTG | tdTomato off-target (short read Illumina MiSeq) |
| SpyCas9 OT4 (chr15) Forward | CACCAGGACCTCTATGGTGC | tdTomato off-target (short read Illumina MiSeq) |
| SpyCas9 OT4 (chr15) Reverse | GCTCCTGCTAGGAGGTATTTGG | tdTomato off-target (short read Illumina MiSeq) |
| SpyCas9 OT5 (chr19) Forward | GGTGCCTCTAAGCATCCTGAAT | tdTomato off-target (short read Illumina MiSeq) |
| SpyCas9 OT5 (chr19) Reverse | AAGTAGCTGGCATGTTCGGT | tdTomato off-target (short read Illumina MiSeq) |
| SauCas9 Forward | GGAAGAGAAATACGTGGCCG | ddPCR cDNA (gene expression) |
| SauCas9 Reverse | GGCTTCTTTCACGTAGTCGC | ddPCR cDNA (gene expression) |
| SauCas9 probe (FAM) | AGAAAGACGGCGAAGTGCGG | ddPCR cDNA (gene expression) |
| EMX1 Forward | GTGTGGTTCCAGAACCGGA | EMX1 on-target (short read Illumina MiSeq) |
| EMX1 Reverse | GCCTGCTTCGTGGCAATG | EMX1 on-target (short read Illumina MiSeq) |

**Supplementary Table 2.**

Assay IDs for qPCR and ddPCR assays.

| Gene Symbol | Assay Catalog # | Manufacturer | Assay/Application |
|---|---|---|---|
| Rbfox3 | PPM60749A | Qiagen | qPCR |
| Gfap | PPM04716A | Qiagen | qPCR |
| Aif1 | PPM03752A | Qiagen | qPCR |
| Itgam | PPM03671F | Qiagen | qPCR |
| Cd19 | PPM03218A | Qiagen | qPCR |
| Sdc1 | PPM03216A | Qiagen | qPCR |
| Cd3e | PPM04598A | Qiagen | qPCR |
| Cd4 | PPM04028F | Qiagen | qPCR |
| Cd8a | PPM04031A | Qiagen | qPCR |
| Il2 | PPM02937C | Qiagen | qPCR |
| Il2ra | PPM03125C | Qiagen | qPCR |
| Il10 | PPM03017C | Qiagen | qPCR |
| Tnf | PPM03113G | Qiagen | qPCR |
| Il1b | PPM03109F | Qiagen | qPCR |
| Ifng | PPM03121A | Qiagen | qPCR |
| Fas | PPM03705B | Qiagen | qPCR |
| Fasl | PPM02926E | Qiagen | qPCR |
| Nfkb1 | PPM02930F | Qiagen | qPCR |
| Il6 | PPM03015A | Qiagen | qPCR |
| Ppih | PPM03699A | Qiagen | qPCR |
| Gapdh | PPM02946E | Qiagen | qPCR |
| MGDC (control) | PPM65836A | Qiagen | qPCR |
| RTC (control) | PPX63340A | Qiagen | qPCR |
| PPC (control) | PPX63339A | Qiagen | qPCR |
| Cdkn1a, 4930567H12Rik | Mm04205640_g1 | ThermoFisher | qPCR |
| Gapdh | Mm99999915_g1 | ThermoFisher | qPCR |
| SpyCas9 NHEJ | dMmuNHS102024435 | Bio-Rad | ddPCR (includes primers and probes (HEX, FAM)) |
| SauCas9 NHEJ | dMmuNHS228153217 | Bio-Rad | ddPCR (includes primers and probes (HEX, FAM)) |

**Supplemental Table 3.**

Off-targets predicted by Cas-OFFinder for SauCas9 (spo4 spacer: 5' TCCAGACATGATAAGATACATTG) evaluated in this study.

| OT | Mismatches | Bulge (Type) | Chromosome (GRCm39) | Sequences (5'→ 3' NNGRRT PAM) |
|---|---|---|---|---|
| 1 | 2 | 2 (RNA) | Chr2 | TCCAG—ATGtTAAGATACATTa TAGAGT<br>TCCAGA—TGtTAAGATACATTa TAGAGT |
| 2 | 2 | 2 (RNA) | Chr18 | TCCAG—ATaATAAaATACATTG AGGAGT<br>TCCAGA—TaATAAaATACATTG AGGAGT<br>TCCAGAtA—ATAAaATACATTG AGGAGT |
| 3 | 3 | 1 (RNA) | Chr1 | TCCtGACATG-TAAGATgCtTTG AGGGGT |
| 4 | 3 | 1 (DNA) | Chr13 | TCaAGACAaGTATAAaATACATTG GTGGGT |
| 5 | 3 | 1 (DNA) | Chr17 | TCaAGACAaGTtTAAGATACATTG GTGGAT<br>TCaAGACAaGtTTAAGATACATTG GTGGAT<br>TCaAGACAaGtTTAAGATACATTG GTGGAT |
| 6 | 3 | 1 (DNA) | Chr4 | TCACAGACATtATAAGAaACATTa CAGAAT |
| 7 | 3 | 1 (RNA) | Chr4 | TCCAtACAT-ATAAGATAgATgG ATGGAT |
| 8 | 3 | 1 (RNA) | Chr9 | TCCAaAaATGATcAGATAC-TTG GTGAAT |
| 9 | 3 | 1 (RNA) | ChrX | T-CAGAaATaAgAAGATACATTG CAGAGT<br>TC-AGAaATaAgAAGATACATTG CAGAGT |
| 10 | 3 | 1 (RNA) | ChrX | TCCAGAC-TcATAAGATACAggG CAGGAT |

**Supplemental Table 4.**

Off-targets predicted by Cas-OFFinder for SpyCas9 (sg298 spacer: 5' AAGTAAAACCTCTACAAATG) evaluated in this study.

| OT | Mismatches | Bulge (Type) | Chromosome (GRCm39) | Sequences (5'→ 3' NGG PAM) |
|---|---|---|---|---|
| 1 | 3 | 0 | Chr4 | AAtTAgAACCTCTACAgATG AGG |
| 2 | 3 | 0 | Chr12 | AAGTAAAACCTCaACAgAaG GGG |
| 3 | 3 | 0 | Chr15 | AAGTAAAACCTCTACcAAaa GGG |
| 4 | 3 | 0 | Chr17 | AAGTAtgACCTaTACAAATG GGG |
| 5 | 3 | 0 | Chr19 | AAtTAAAAgCTCTACAAAaG GGG |

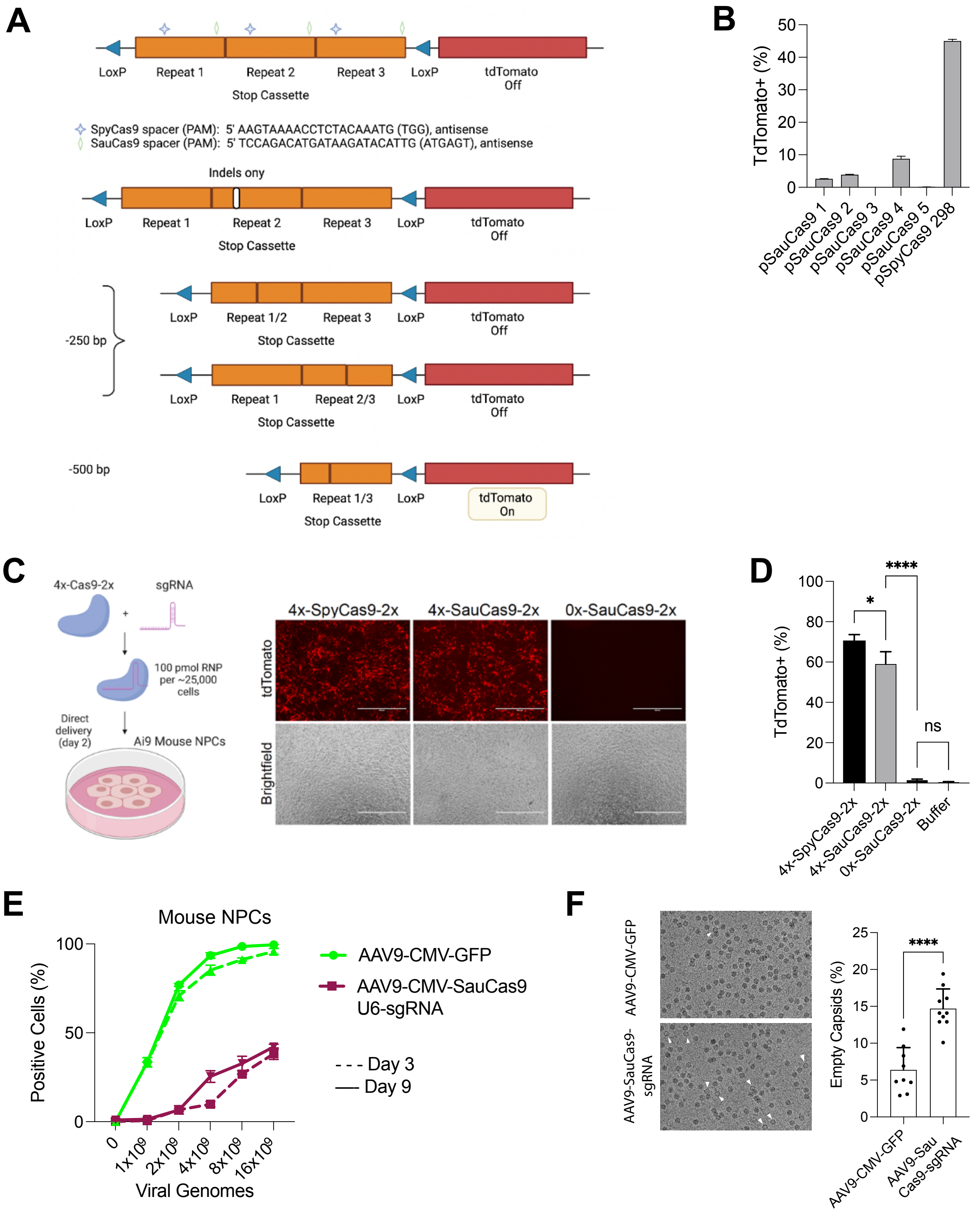

A
LoxP
Repeat 1
Repeat 2
Repeat 3
LoxP
tdTomato Off
Stop Cassette
SpyCas9 spacer (PAM): 5' AAGTAAAACCTCTACAAATG (TGG), antisense
SauCas9 spacer (PAM): 5' TCCAGACATGATAAGATACATTG (ATGAGT), antisense
Indels ony
-250 bp
Repeat 1/2
Repeat 2/3
-500 bp
Repeat 1/3
tdTomato On
B
TdTomato+ (%)
pSauCas9 1
pSauCas9 2
pSauCas9 3
pSauCas9 4
pSauCas9 5
pSpyCas9 298
C
4x-Cas9-2x
sgRNA
100 pmol RNP per ~25,000 cells
Direct delivery (day 2)
Ai9 Mouse NPCs
4x-SpyCas9-2x
4x-SauCas9-2x
0x-SauCas9-2x
tdTomato
Brightfield
D
TdTomato+ (%)
*
****
ns
4x-SpyCas9-2x
4x-SauCas9-2x
0x-SauCas9-2x
Buffer
E
Mouse NPCs
Positive Cells (%)
Viral Genomes
0
1x10^9
2x10^9
4x10^9
8x10^9
16x10^9
AAV9-CMV-GFP
AAV9-CMV-SauCas9 U6-sgRNA
Day 3
Day 9
F
AAV9-CMV-GFP
AAV9-SauCas9-sgRNA
Empty Capsids (%)
****
AAV9-CMV-GFP
AAV9-Sau Cas9-sgRNA

**Supplementary Figure 1. Adapting tdTomato reporter system for SauCas9 genome editing with RNPs and AAVs in vitro.**

A) Schematic of tdTomato locus in Ai9 mice, created with Biorender.com. Three stop repeats are flanked by LoxP sites upstream of the tdTomato reporter, which separates tdTomato from its promoter within the Rosa26 locus. The relative target sites for SpyCas9 (sg298) and SauCas9 (spacer4) are shown including the spacer and PAM sequences. After dsDNA breaks, there can be several editing outcomes, including indels only in any of the three repeats (one shown here), deletions of approximately 250bp from neighboring repeats, or large deletions of approximately 500bp from distal repeats, which is the only outcome sufficient to turn on the tdTomato reporter. Therefore, tdTomato fluorescent signal is an underestimate of total editing events.

B) Flow cytometry results from vitro experiment using neural precursor cells (NPCS) from E13.5 Ai9 mice to measure editing after nucleofection with six plasmids: five plasmids encoded SauCas9 with five different spacer sequences and one plasmid encoded SpyCas9 with sg298 as a positive control. Puromycin selection was not performed. Out of the five SauCas9 guides spacer 4 performed the best and was selected for further study.

C) Schematic of RNP formation and direct delivery in Ai9 tdTomato mouse NPCs, created with BioRender.com. Representative fluorescent micrographs showing tdTomato signal in mouse NPCs five-days after treatment with RNPs prior to flow cytometry. Scale bar: 400μm.

D) Flow cytometry results of editing as measured by tdTomato expression after cell penetrant Cas9 RNPs were added to mouse neural precursor cells (NPCs) via “direct delivery” into the cell culture supernatant (100pmol, final 1uM concentration). Treatments were performed in triplicate and statistical significance was measured by one-way ANOVA with multiple comparisons (**** $p<0.001$, * $p<0.05$, ns = not significant).

E) Flow cytometry results after viral transduction of mouse NPCs with a virus encoding the GFP transgene neared 100% when 10,000 cells received 2e9 viral genomes (vg), while editing as measured by tdTomato expression reached a maximum of approx. 40% at the highest tested dose of 16e9 vg. Above 8e9 vg, cell viability was visibly decreased.

F) Cryo-electron microscopy of AAV9 capsids. Three individuals blinded to the identify of 9-10 images per group counted empty (arrows) versus full capsids in ImageJ. The three counts per image were averaged then plotted between the two groups (student’s t-test, **** $p<0.001$).

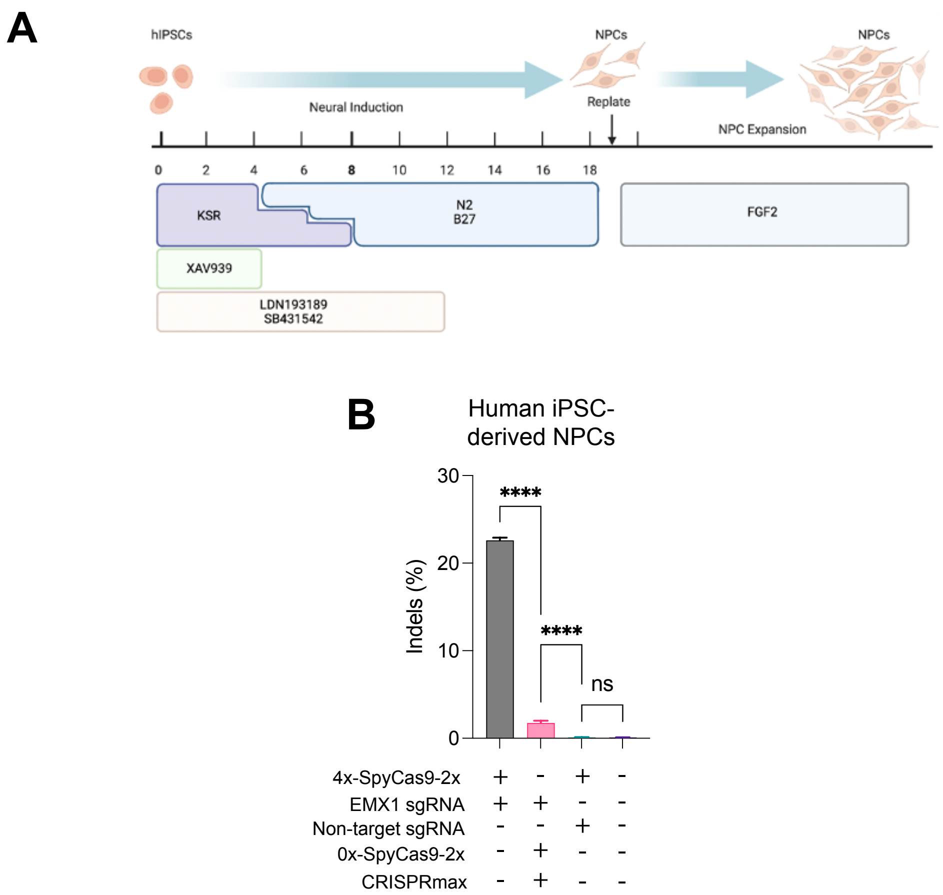


**Supplemental Figure 2. Direct delivery of 4xNLS-Cas9 RNPs results in significant editing of primary human NPCs.**

A) Schmatic of human induced pluripotent stem cell (iPSC) differentiation into neural precursor cells (NPCs) and expansion. Created with Biorender.com.

B) Quantification of editing of EMX1 locus in iPSC-derived NPCs with direct delivery addition of SpyCas9 protein, targeting and non-targeting guide RNAs,and CRISPRmax transfection reagent (n=3 replicates, one-way ANOVA, ****p<0.001).

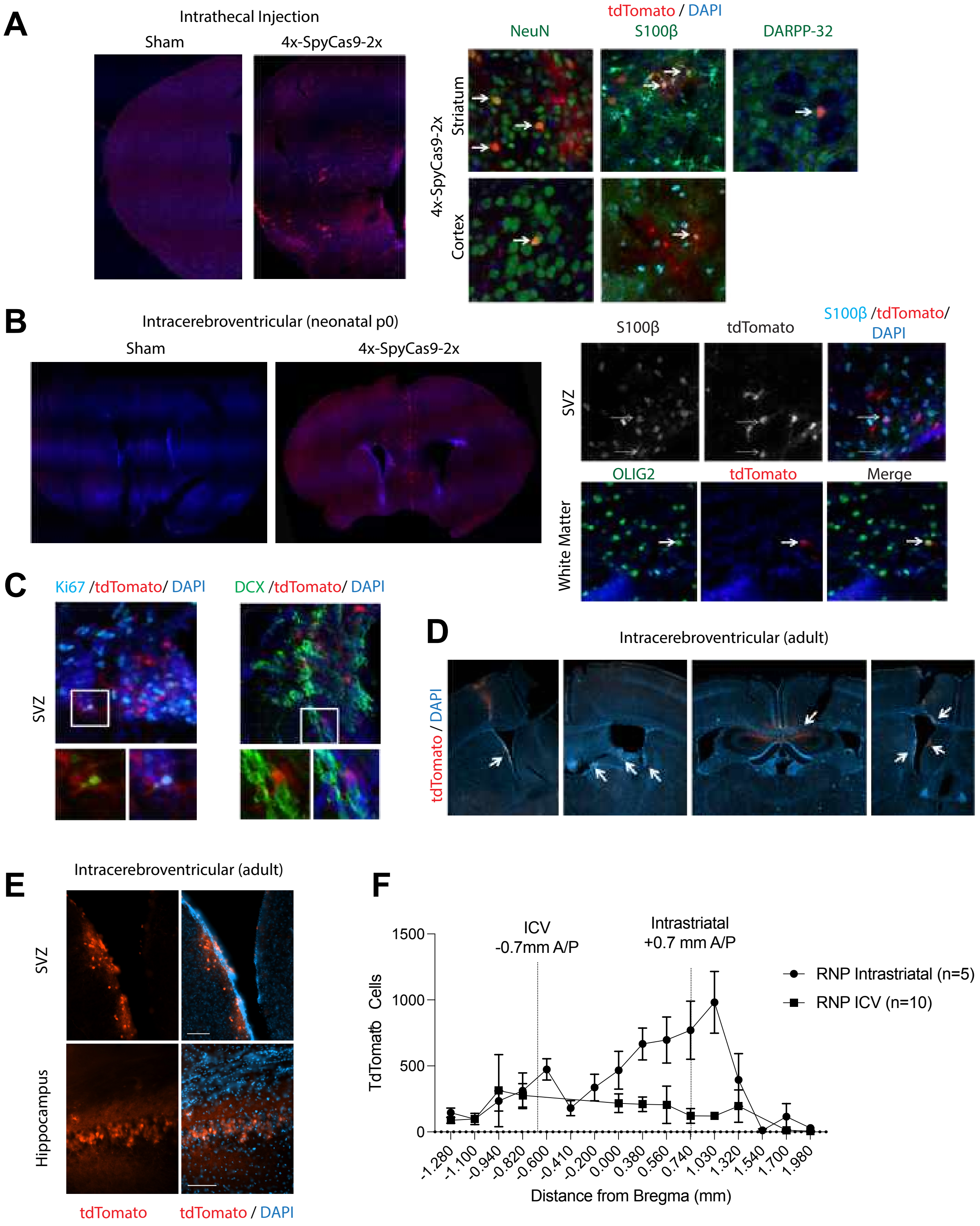

A
Intrathecal Injection
Sham
4x-SpyCas9-2x
tdTomato / DAPI
NeuN
S100β
DARPP-32
4x-SpyCas9-2x
Striatum
Cortex
B
Intracerebroventricular (neonatal p0)
Sham
4x-SpyCas9-2x
S100β
tdTomato
S100β /tdTomato/ DAPI
SVZ
OLIG2
tdTomato
Merge
White Matter
C
Ki67 /tdTomato/ DAPI
DCX /tdTomato/ DAPI
SVZ
D
Intracerebroventricular (adult)
tdTomato / DAPI
E
Intracerebroventricular (adult)
SVZ
Hippocampus
tdTomato
tdTomato / DAPI
F
ICV
-0.7mm A/P
Intrastriatal
+0.7 mm A/P
RNP Intrastriatal (n=5)
RNP ICV (n=10)
TdTomato Cells
1500
1000
500
0
-1.280
-1.100
-0.940
-0.820
-0.600
-0.410
-0.200
0.000
0.380
0.560
0.740
1.030
1.320
1.540
1.700
1.980
Distance from Bregma (mm)

**Supplemental Figure 3. Intrathecal and intracerebroventricular (ICV) administration of Cas9 RNPs.**

A) Intrathecal injection of 300µM 4x-SpyCas9-2x into the sub-arachnoid space in adult Ai9 mice analyzed 21-days post-injection. Edited NeuN+ neurons and S100beta+ glial cells were detected in the cortex and striatum of one mouse, including DARPP-32 medium spiny neurons. Only 1 of 3 mice at the 300µM dose showed tdTomato signal in the brain. Additional mice that received intrathecal injection of RNP at 100µM (n=4), 200µM (n=4), and 400µM (n=5) did not have detectable editing in the brain or spinal cord.

B) ICV injection of 100µM 4x-SpyCas9-2x into the lateral ventricle of neonatal p0 mice analyzed 21-days post-injection. Edited S100beta+ and OLIG2+ glial cells were identified in the subventricular zone (SVZ) and white matter. Representative image shown (n=1 of 5). All 5 injected mice had similar levels of editing, however, some also showed signs of ventriculomegaly.

C) ICV injection of p0 mice also led to tdTomato expression (editing) in neural stem/progenitor cells that express Ki67 and DCX.

D) ICV injection of adult mice with 100-400µM 4x-SpyCas9-2x led to edited tdTomato+ cells in the SVZ, choiroid plexus, ventricle, corpus callosum, and hippocampus (images shown from 3 different biological replicates).

E) Zoomed (20x magnification) image of tiled slide scan of adult ICV injection in two replicates showing tdTomato+ cells in hippocampus and sub-ventricular zone (SVZ). Scale bar: 100µm.

F) Quantification of tdTomato+ cells in adult Ai9 mice for ICV route versus intrastriatal injections. From n=5 injections, the intrastriatal route resulted in an average of 3631 tdTomato+ cells per striatum (range 1943 to 6209).  From n=10 injections, all 10 mice had tdTomato signal in the tissue near the ventricles, averaging 859 tdTomato+ cells per brain (range 295 to 4329), demonstrating direct injection of RNP is most effective for editing in the mouse brain.

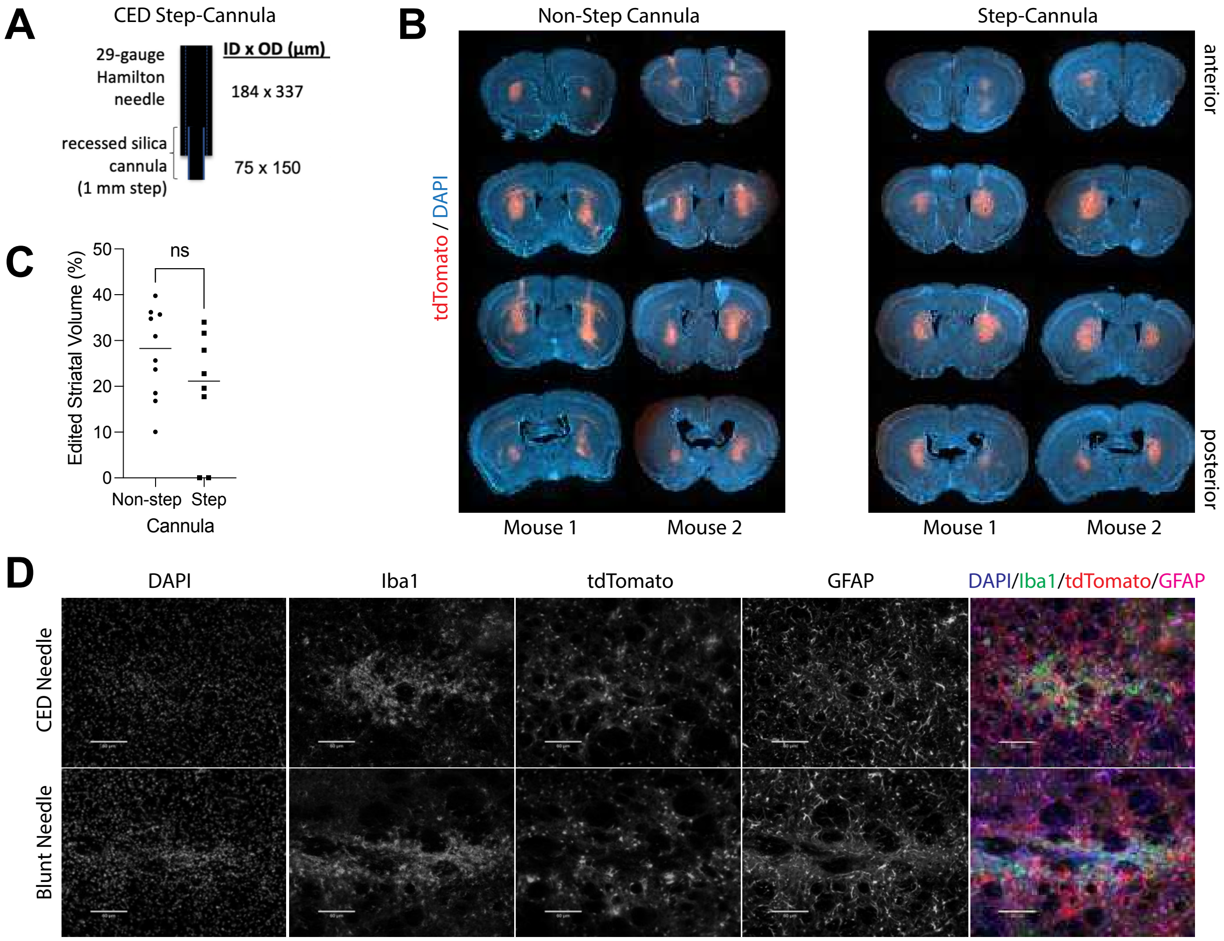


**Supplemental Figure 4. Convection enhanced delivery of RNPs.**

A) Schematic of convection enhanced delivery (CED) needle with step-cannula.

B) Representative images of two biological replicates of intrastriatal bilateral injections of 4x-SpyCas9-2x RNP at 25µM (125pmol) per hemisphere at 7-days post-injection.

C) Quantification of edited striatal volume in the non-step cannula (n=10 techncial replicates) and the step-cannula (n=8 technical replicates) showed no signficiant difference. Of note, the step-cannula group contained two failed injections (along x-axis). Additionally, two mice did not recover from surgery in the step-cannula group and one mouse did not recover from surgery in the non-step cannula group (data not shown).

D) Representative images of Iba1 and GFAP staining near the injeciton site in the striatum demonstrated similar levels of microglial and glial cell reaction between the step and non-step cannulas. Scale bar: 50µm.

**A**

| | Theoretical Charge | Elution KCl (mM) |
|---|---|---|
| 0x-SpyCas9-2x | + 30 | 470 |
| 4x-SpyCas9-2x | + 50 | 690 |
| 0x-SauCas9-2x | + 40 | 660 |
| 4x-SauCas9-2x | + 61 | 760 |
| 3x-SauCas9-2x | + 56 | 730 |
| 2x-SauCas9-2x | + 51 | 700 |

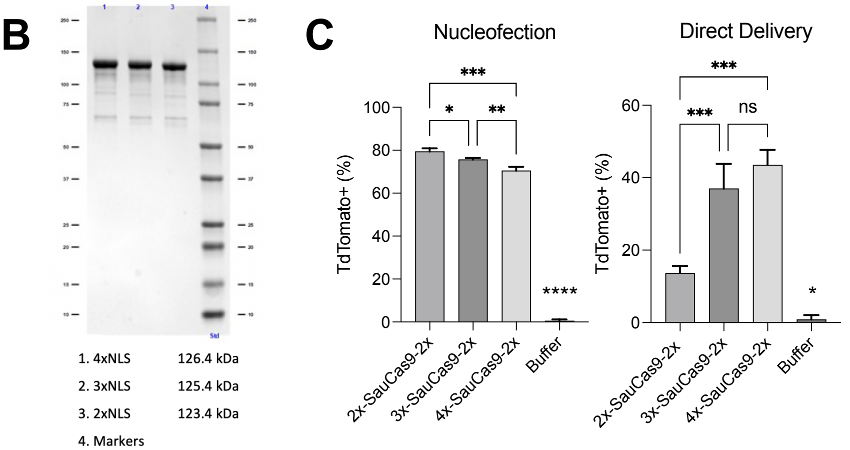


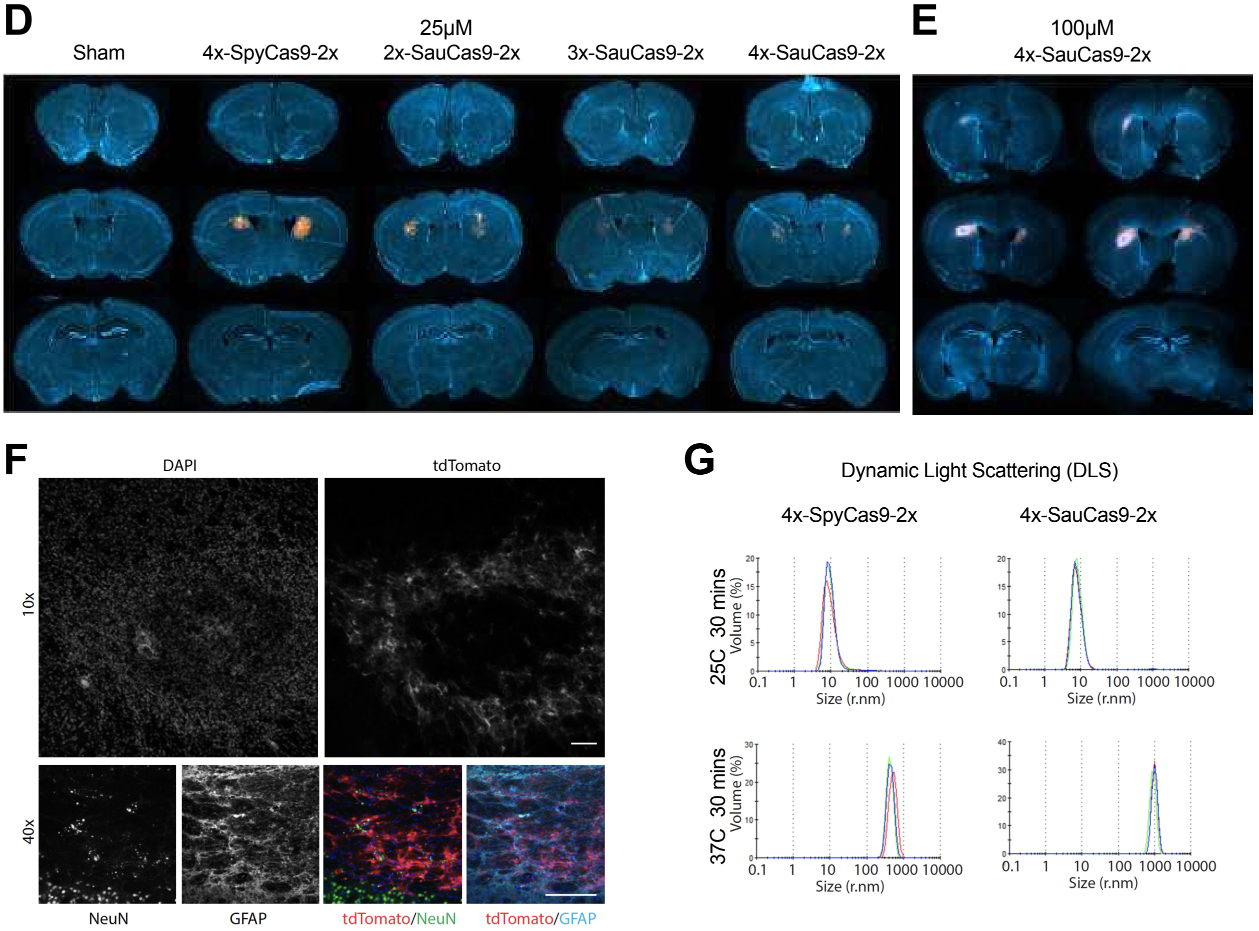

**Supplemental Figure 5. Optimizing SauCas9 RNP NLS structure for self-delivery in vivo.**

A) Theoretical charge and measured elution concentration of SpyCas9 and SauCas9 variants with different SV40 NLS architectures on the N and C terminus.

B) SDS-PAGE gel showing the expression purity and size of the three SauCas9 NLS variants.

C) Flow cytometry results of editing in Ai9 NPCs from three SauCas9 variants with 2, 3, and 4 copies of SV40 NLS on the N-terminus. The 4x-SauCas9-2x variant was most effective at direct entry into the cell (added to cell culture supernatant at 100pmol, 1μM final concentration, 48-hours after seeding the cells), however the 2x-SauCas9-2x variant was most effective at editing the cells when nucleofected, suggesting the NLS can interfere with editing function but enhances cell entry.

D) Representative images (1 of 2 biological replicates shown) of bilateral intrastriatal injection of RNPs (25uM, 125 pmol dose) analyzed at 21-days shows very limited editing with SauCas9 RNPs in vivo. All groups lagged behind 4x-SpyCas9-2x RNP at in vivo efficiency (20μm thick OCT- embedded sections).

E) Representative images (2 of 3 biological replicates) of bilateral intrastriatal injection of RNPs (100μM, 500 pmol dose) analyzed at 21-days shows very limited editing with SauCas9 RNPs in vivo (50μm thick agarose embedded sections).

F) Confocal images of edited striatum in 4x-SauCas9-2x RNP group 21-days after injection at 500pmol. The tdTomato often had a “donut shape” pattern where signal was lost from the center, concomitant with loss of NeuN and enhanced GFAP staining, suggesting dose-limiting toxicity. 10x images and 40x images were imaged on different days and reflect the same group, but not the same section. Scale bar: 100 μm.

G) Dynamic light scattering experiment with Cas9 RNPs incubated at 25 or 37C showing increase in size distribution by mass peak over time. Larger RNPs could indicate aggregation, likely following temperature-induced protein misfolding.

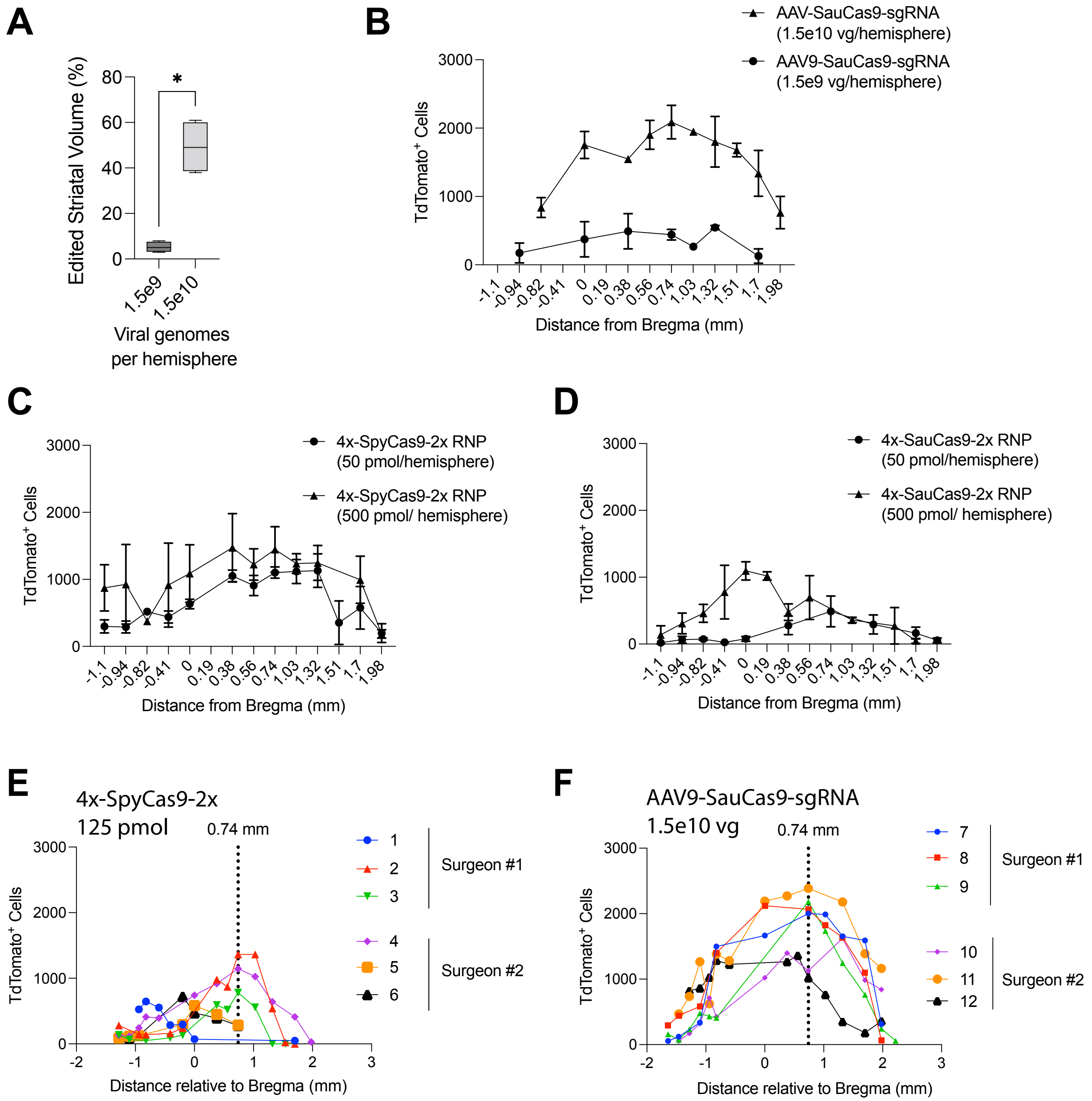

A
Edited Striatal Volume (%)
80
60
40
20
0
*
1.5e9
1.5e10
Viral genomes per hemisphere
B
AAV-SauCas9-sgRNA (1.5e10 vg/hemisphere)
AAV9-SauCas9-sgRNA (1.5e9 vg/hemisphere)
TdTomato+ Cells
3000
2000
1000
0
-1.1 -0.94 -0.82 -0.41 0 0.19 0.38 0.56 0.74 1.03 1.32 1.51 1.7 1.98
Distance from Bregma (mm)
C
4x-SpyCas9-2x RNP (50 pmol/hemisphere)
4x-SpyCas9-2x RNP (500 pmol/ hemisphere)
TdTomato+ Cells
Distance from Bregma (mm)
D
4x-SauCas9-2x RNP (50 pmol/hemisphere)
4x-SauCas9-2x RNP (500 pmol/ hemisphere)
TdTomato+ Cells
Distance from Bregma (mm)
E
4x-SpyCas9-2x
125 pmol
0.74 mm
1
2
3
Surgeon #1
4
5
6
Surgeon #2
TdTomato+ Cells
-2 -1 0 1 2 3
Distance relative to Bregma (mm)
F
AAV9-SauCas9-sgRNA
1.5e10 vg
0.74 mm
7
8
9
Surgeon #1
10
11
12
Surgeon #2
TdTomato+ Cells
Distance relative to Bregma (mm)

**Supplemental Figure 6. Dose dependent effects on distribution of Cas9-AAV and Cas9-RNPs in the brain.**

A) Edited tissue volume calculated as a percentage of total striatum (%) for AAV9-CMV-SauCas9-U6-sgRNA injected at 3e8 vg/μL and 3e9 vg/μL (1.5e9-1.5e10 vg/hemisphere) analyzed at 90-days post-injection. Unpaired t-test, *$p<0.05$.

B) Total tdTomato+ cell counts per hemisphere (all brain regions) along the rostral-caudal axis for AAV9-CMV-SauCas9-U6-sgRNA injected at 3e8 vg/μL and 3e9 vg/μL (1.5e9-1.5e10 vg/hemisphere) analyzed at 90-days post-injection. Mean +/- standard error (n=4 injections).

C) Total tdTomato+ cell counts per hemisphere (all brain regions) along the rostral-caudal axis for 4x-SpyCas9-2x RNPs at 50 or 500 pmol dose analyzed up to 90-days post-injection. Mean +/- standard error (n=4 injections).

D) Total tdTomato+ cell counts per hemisphere (all brain regions) along the rostral-caudal axis for 4x-SauCas9-2x RNPs at 50 or 500 pmol dose analyzed up to 90-days post-injection. Mean +/- standard error (n=4 injections).

E) Individual replicates of total tdTomato+ cell counts per hemisphere (all brain regions) of mice that received injections of 4x-SpyCas9-2x analyzed at 21-days. Cas9 RNPs (25μM, 125pmol, n=6) had a maximum number of 1000 edited cells near the injection site (dotted line 0.74 mm relative to Bregma).

F) Individual replicates of total tdTomato+ cell counts per hemisphere (all brain regions) of mice injected with AAV9-CMV-SauCas9-U6-sgRNA analyzed at 21-days. Cas9 AAVs (3e9 vg/μL,1.5e10 vg/hemisphere, n=6) resulted in a maximum of 2000 edited cells near the injection site and diffused more broadly along the anterior-posterior axis.

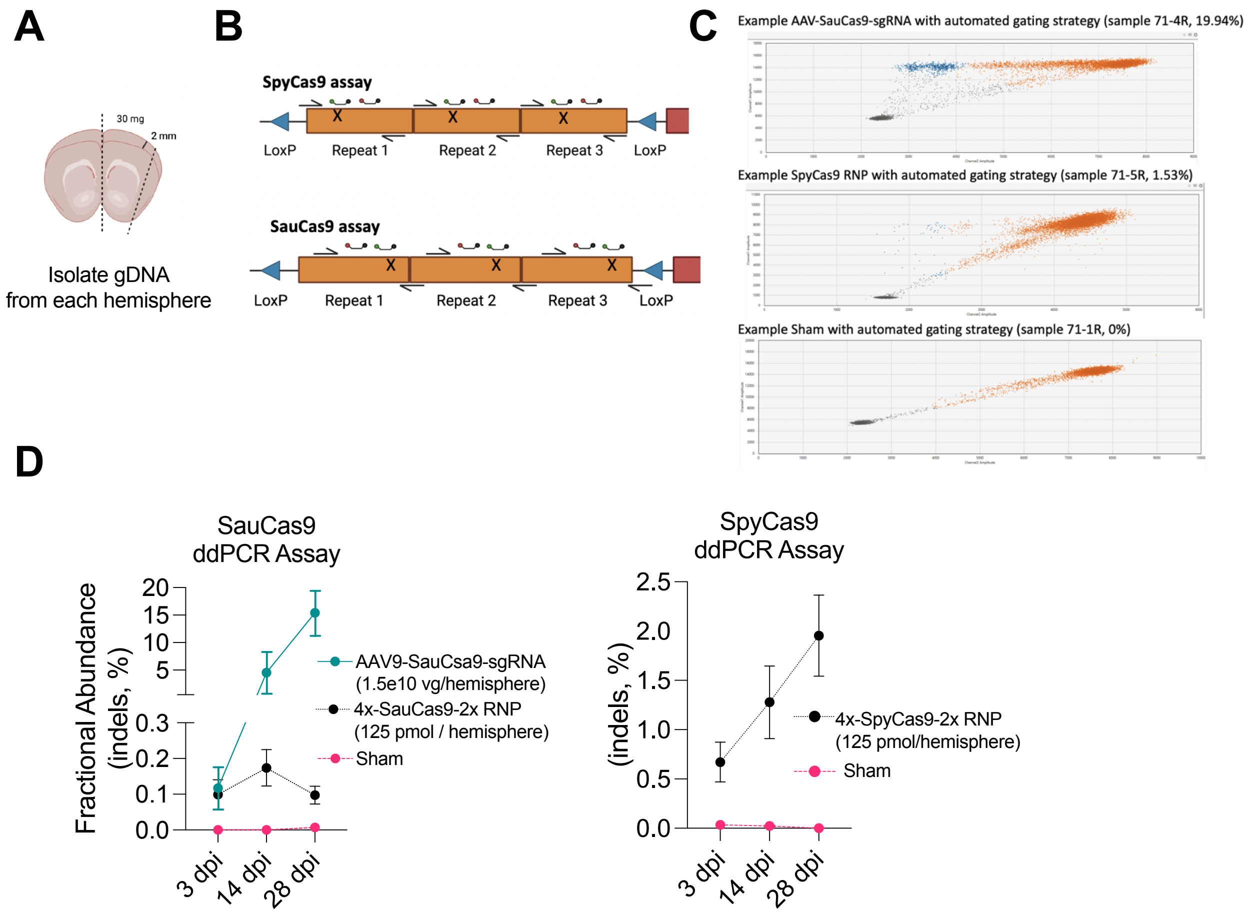


**Supplemental Figure 7. Development of ddPCR assay for editing outcomes with SpyCas9 -sg298 and SauCas9-spacer4 at tdTomato locus**.

A) Schematic of tissue collection. Approximately 30 mg of brain tissue (2 mm thick) was isolated from one hemisphere of the mouse striatum, encompassing the intended injection site. Tissue was flash frozen and later processed for genomic DNA (gDNA). Image made using Biorender.com.

B) Schematic of PCR products, approximately 160bp within each of the three stop repeats containing the cut site. Two probes bind to the amplicon. The FAM-labeled probe is the reference, while the HEX-labeled probe sits directly over the intended cut-site and will be lost due to indels or large deletions. Due to different gRNA requirements, each ortholog was assessed with a different primer/probe set. Droplets that loses HEX in greater proportion to FAM indicate an editing event repaired with NHEJ/MMEJ resulting in indels.

C) Representative results showing FAM+ / HEX – population (blue) in Cas9-AAV, RNP, and sham injected mice, signifying disruption of the tdTomato locus at the cut site.

D) Quantification of digital droplet PCR (ddPCR) assays show detectable editing in AAV9-SauCas9-sgRNA, 4x- SauCas9-2x NLS, and 4x-SpyCas9-2x groups (25µM, 125pmol, n=6 technical replicates) in 30mg tissue hemispheres (2-mm thick) above background (sham).

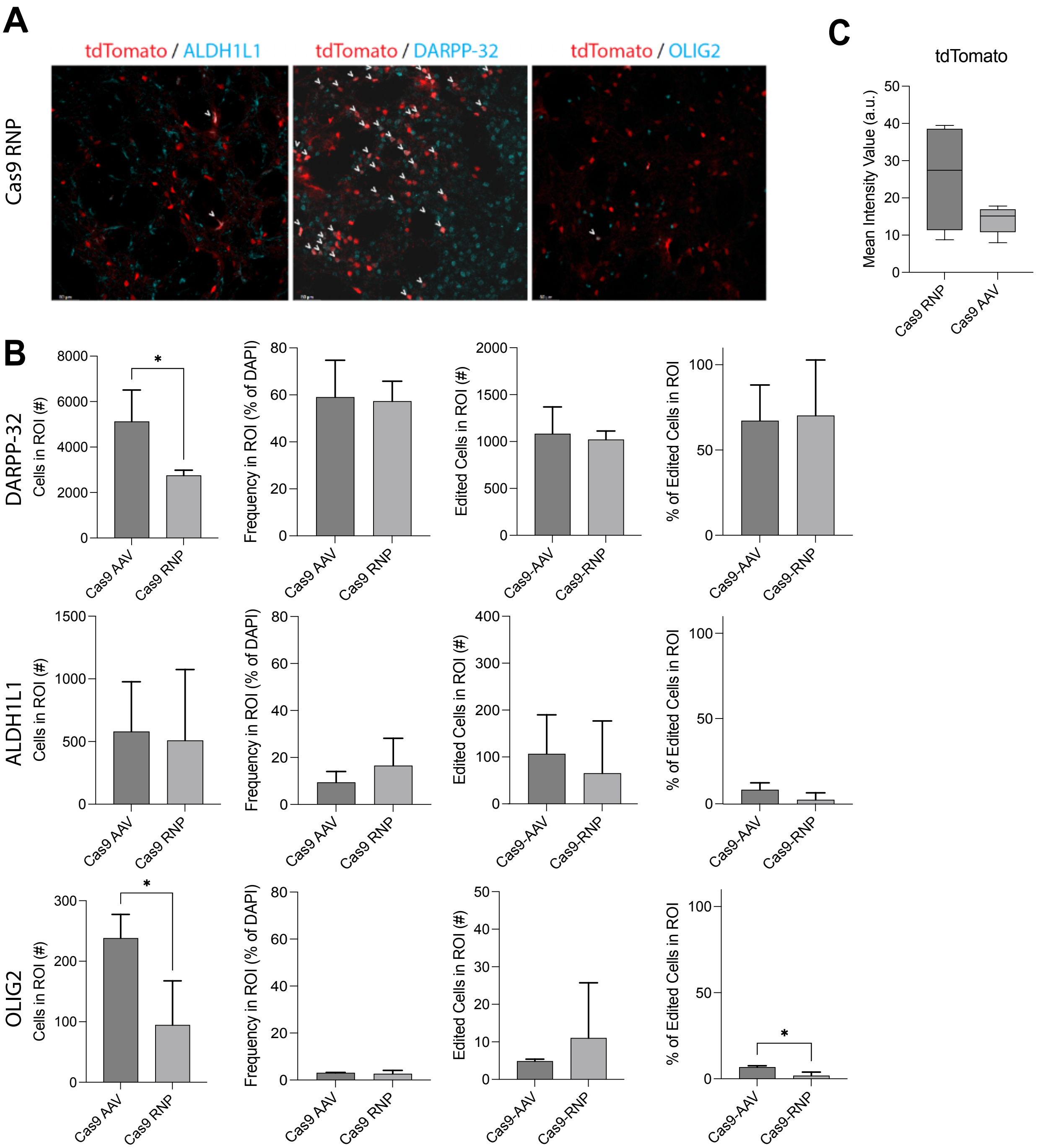

A
tdTomato / ALDH1L1
tdTomato / DARPP-32
tdTomato / OLIG2
Cas9 RNP
C
tdTomato
Mean Intensity Value (a.u.)
Cas9 RNP
Cas9 AAV
B
DARPP-32
ALDH1L1
OLIG2
Cells in ROI (#)
Frequency in ROI (% of DAPI)
Edited Cells in ROI (#)
% of Edited Cells in ROI
Cas9 AAV
Cas9 RNP
Cas9-AAV
Cas9-RNP
*

**Supplemental Figure 8. Edited Cell Distribution and tdTomato intensity.**

A) Representative images of ALDH1L1, DARPP-32, and OLIG2 immunostaining staining (cyan) with tdTomato (red) in the mouse hippocampus 21-days post-delivery of Cas9 RNPs (25µM). Arrows indicate cells co-expressing tdTomato and each cell marker. Scale bar: 50 µm.

B) Quantification of cell type editing for DARPP-32 (medium spiny neurons), ALDH1L1 (astrocytes), and OLIG2 (oligodendrocytes). Graphs in the first column show the total number of cells expressing the indicated marker within the region of interest (ROI). ROIs were often larger for Cas9-AAV due to greater lateral diffusion and thus contain more cells (n=3 injections, unpaired t-test, * $p<0.05$). The second column shows the frequency of each cell type in the ROI as a percentage of total cells. There was no significant difference in the frequency of each cell type within the ROI (i.e., no population death or expansion in response to treatment), although there was a trend towards more ALDH1L1+ cells in the Cas9 RNP group. Column 3 shows the number of edited (tdTomato+) cells within the ROI expressing the given marker. Despite larger ROIs, there was no significant increase in edited cell counts in the Cas9 RNP group. The fourth column shows the frequency of edited cells within the ROI. In both Cas9 AAV and RNP, edited DARPP-32+ cells accounted for greater than half of the total edited cells. The DARPP-32 data complements the NeuN data in Figure 1J, demonstrating that Cas9 RNPs result in the same or more edited neurons than Cas9 AAVs within a given area. Glial cells accounted for a significantly greater proportion of total edited cells in the Cas9 AAV group compared to the Cas9 RNP group (* $p<0.05$).

C) Per cell mean intensity values of tdTomato were quantified within the ROIs for Cas9 AAV and RNPs. The tdTomato signal was often brighter in the Cas9 RNP group but was not statistically different (n=4-6 injections, unpaired t-test, ns). This observation could perhaps be due to differences in guide RNA editing outcomes or a greater frequency of biallelic editing in the Cas9-RNP group as tdTomato signal on requires two simultaneous cuts. Cas expression from AAV genome likely has a greater lag time to produce multiple Cas9 molecules required for gene excision and could thus lead to greater number of indels not captured by fluorescence measurements.

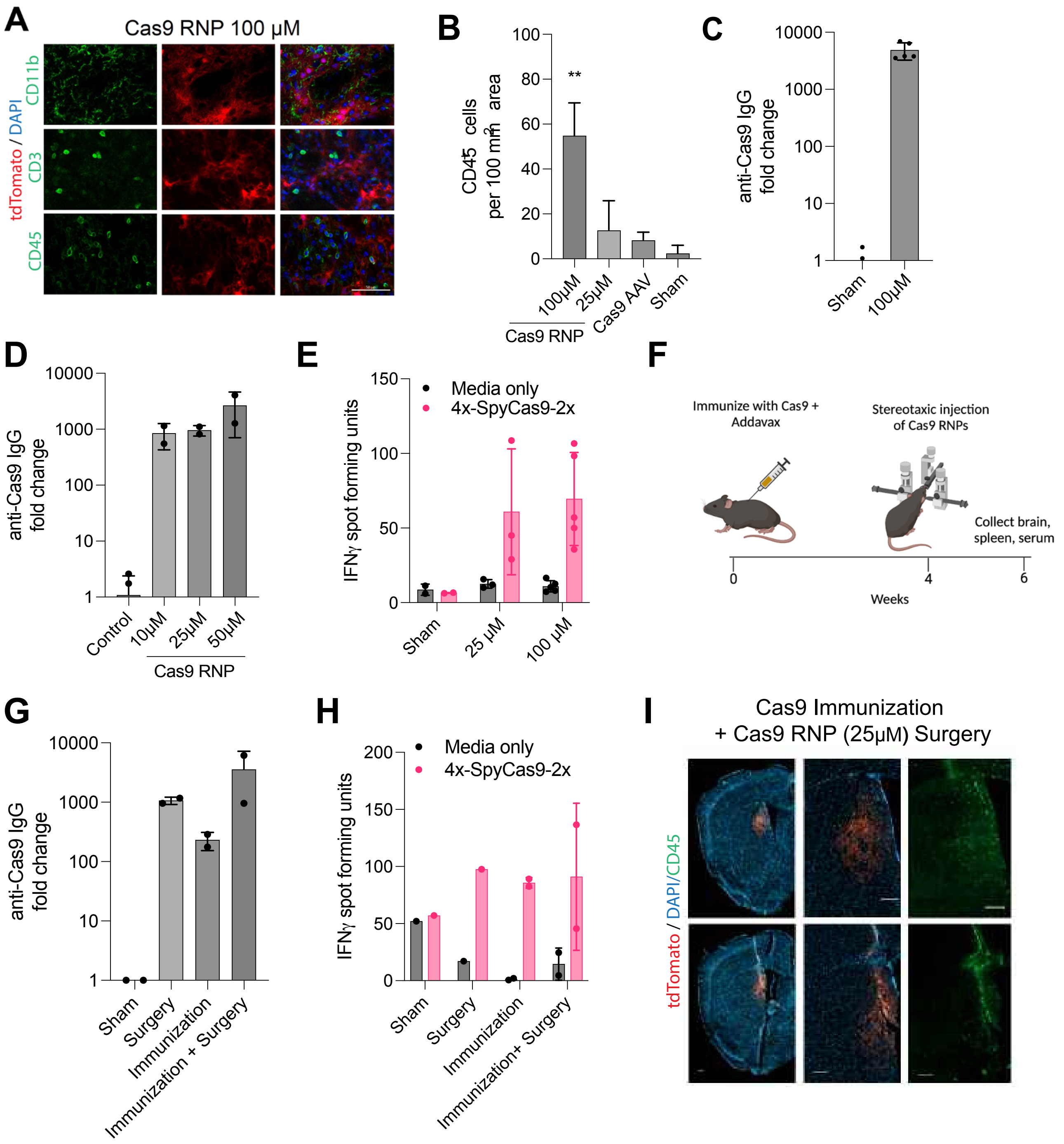

A
Cas9 RNP 100 µM
tdTomato / DAPI
CD11b
CD3
CD45
B
CD45+ cells per 100 mm² area
100
80
60
40
20
0
**
100µM
25µM
Cas9 AAV
Sham
Cas9 RNP
C
anti-Cas9 IgG fold change
10000
1000
100
10
1
Sham
100µM
D
anti-Cas9 IgG fold change
Control
10µM
25µM
50µM
Cas9 RNP
E
IFNγ spot forming units
150
100
50
0
Media only
4x-SpyCas9-2x
Sham
25 µM
100 µM
F
Immunize with Cas9 + Addavax
Stereotaxic injection of Cas9 RNPs
Collect brain, spleen, serum
0
4
6
Weeks
G
anti-Cas9 IgG fold change
Sham
Surgery
Immunization
Immunization + Surgery
H
IFNγ spot forming units
200
150
100
50
0
Media only
4x-SpyCas9-2x
Sham
Surgery
Immunization
Immunization+ Surgery
I
Cas9 Immunization
+ Cas9 RNP (25µM) Surgery
tdTomato / DAPI/CD45

**Supplemental Figure 9. Cellular and humoral response to Cas9 RNPs at additional doses and following immunization with Addavax with Cas9 protein.**

A) Representative staining of CD45, CD3, and CD11b in the Cas9 RNP 500pmol dose at 21-days. Scale bar: 50µm.

B) Quantification of CD45 immune cells at 21-days in Cas9 RNP 100 µM (500 pmol) dose compared to Cas9 RNP 25 µM (125 pmol), Cas9 AAV, and sham (as shown in Figure 3C, oneway ANOVA, ** $p < 0.01$).

C) Anti-Cas9 IgG measured by ELISA from serum collected 21-days post-treatment with Cas9 RNP 100 µM dose had similar fold change as the Cas9 RNP 25 µM dose (Figure 3E).

D) Anti-Cas9 IgG measured by ELISA from serum collected 90-days post-treatment with Cas9 RNP at 10 µM, 25 µM, and 100 µM doses.

E) Anti-Cas9 cellular response (interferon-gamma spot forming units) measured by ELISpot from splenocytes collected 21-days post-treatment with Cas9 RNP at 25 µM and100 µM compared to sham mice, restimulated in culture for 48 hours with media only or 4x-SpyCas9-2x.

F) Schematic of immunization strategy (25µg 4x-SpyCas9-2x injected subcutaneously with AddaVax, followed by stereotaxic surgery with 25 µM (10µL at 4.15 µg/µL: 41.5 µg) 4x-SpyCas9-2x). Created with Biorender.com.

G) Anti-Cas9 IgG measured by ELISA from serum collected 6-weeks post-immunization (2 weeks post-surgery if applicable). Sham mice received Addavax alone and no surgery, immunized mice received Addavax with 4x-SpyCas9-2x protein subcutaneously only, and surgery mice received25 µM 4x-SpyCas9-2x RNP into the striatum.

H) Anti-Cas9 cellular response (interferon-gamma spot forming units) measured by ELISpot from splenocytes collected 6 weeks post-immunization (2 weeks post-surgery if applicable).

I) Representative image of tdTomato editioning in the brain at 6 weeks post-immunization (2 weeks post-surgery) in a mouse that received both immunization and surgery. Many CD45+ cells were observed near tdTomato+ cells and along needle injection track, which resembled the response to higher doses of Cas9-RNPs. Edited cells persisted to the latest measured time point of 2-weeks. Scale bar: 200 µm.

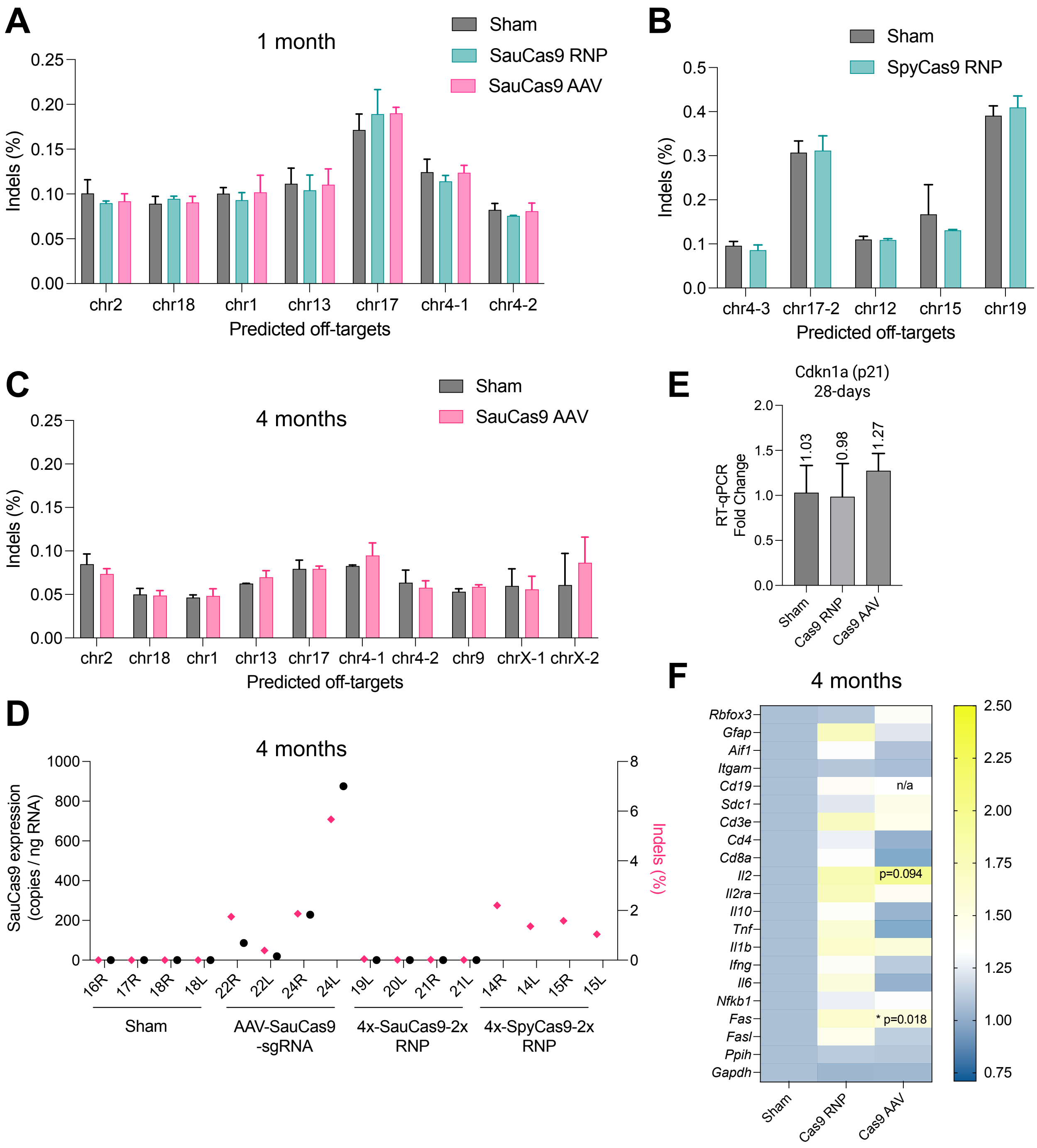
A
1 month
Sham
SauCas9 RNP
SauCas9 AAV
Indels (%)
0.25
0.20
0.15
0.10
0.05
0.00
chr2
chr18
chr1
chr13
chr17
chr4-1
chr4-2
Predicted off-targets
B
Sham
SpyCas9 RNP
Indels (%)
0.5
0.4
0.3
0.2
0.1
0.0
chr4-3
chr17-2
chr12
chr15
chr19
Predicted off-targets
C
4 months
Sham
SauCas9 AAV
Indels (%)
chr2
chr18
chr1
chr13
chr17
chr4-1
chr4-2
chr9
chrX-1
chrX-2
Predicted off-targets
E
Cdkn1a (p21)
28-days
RT-qPCR
Fold Change
2.0
1.5
1.0
0.5
0.0
1.03
0.98
1.27
Sham
Cas9 RNP
Cas9 AAV
D
4 months
SauCas9 expression
(copies / ng RNA)
1000
800
600
400
200
0
8
6
4
2
0
Indels (%)
16R
17R
18R
18L
22R
22L
24R
24L
19L
20L
21R
21L
14R
14L
15R
15L
Sham
AAV-SauCas9
-sgRNA
4x-SauCas9-2x
RNP
4x-SpyCas9-2x
RNP
F
4 months
Rbfox3
Gfap
Aif1
Itgam
Cd19
Sdc1
Cd3e
Cd4
Cd8a
Il2
Il2ra
Il10
Tnf
Il1b
Ifng
Il6
Nfkb1
Fas
Fasl
Ppih
Gapdh
n/a
p=0.094
* p=0.018
2.50
2.25
2.00
1.75
1.50
1.25
1.00
0.75
Sham
Cas9 RNP
Cas9 AAV

**Supplemental Figure 10. Assessment of SauCas9 transgene expression and off-target editing *in vivo*.**

A) Next-generation sequencing (NGS) results evaluating indels at seven selected sites predicted as potential off-targets by Cas-OFFinder for SauCas9 RNP, AAV, and sham treated mice 28-days post-injection. No significant differences were found.

B) NGS results evaluating indels at five selected sites predicted by Cas-OFFinder for SauCas9 RNP and sham treated mice 28-days post-injection. No significant differences were found.

C) NGS results evaluating indels at ten selected sites predicted as potential off-targets by Cas-OFFinder for SauCas9 AAV and sham treated mice 4-months post-injection. No significant differences were found.

D) Digital droplet PCR assay for on-target indels at 4-months post-treatment (pink diamonds, right y-axis, maximum 2.2% in SpyCas9 RNP and 5.67% in SauCas9 AAV). Of note, a different lot of AAV9-SauCas9-sgRNA was in the 1-month experiments (Supplemental Figure 5G). SauCas9 expression was detected 4-months post-treatment only in the Cas9-AAV group (black circles, left y-axis, maximum 875 copies/ng), which correlated with indels. SauCas9 expression was not evaluated in the SpyCas9-RNP group. The result of the ddPCR (copies/µL) was multiplied by the total ddPCR reaction volume and divided by the cDNA input volume to calculate cDNA copies/µL. Then values were multiplied by the cDNA dilution factor, multiplied by the reverse transcription reaction volume, and divided by the starting amount of RNA (500 ng) to calculate copies/ng RNA.

E) No significant activation of p21 (Cdkn1a) was found via RT-qPCR at 28-days post-treatment with Cas9 RNP or Cas9 AAV.

F) Gene expression heat map showed significant upregulation of Fas in the Cas9 AAV group 4 months post-treatment with Cas9 AAV (n=4, unpaired t-test compared to sham, $*p<0.05$). Cd19 was not detected in the Cas9-AAV group.

A

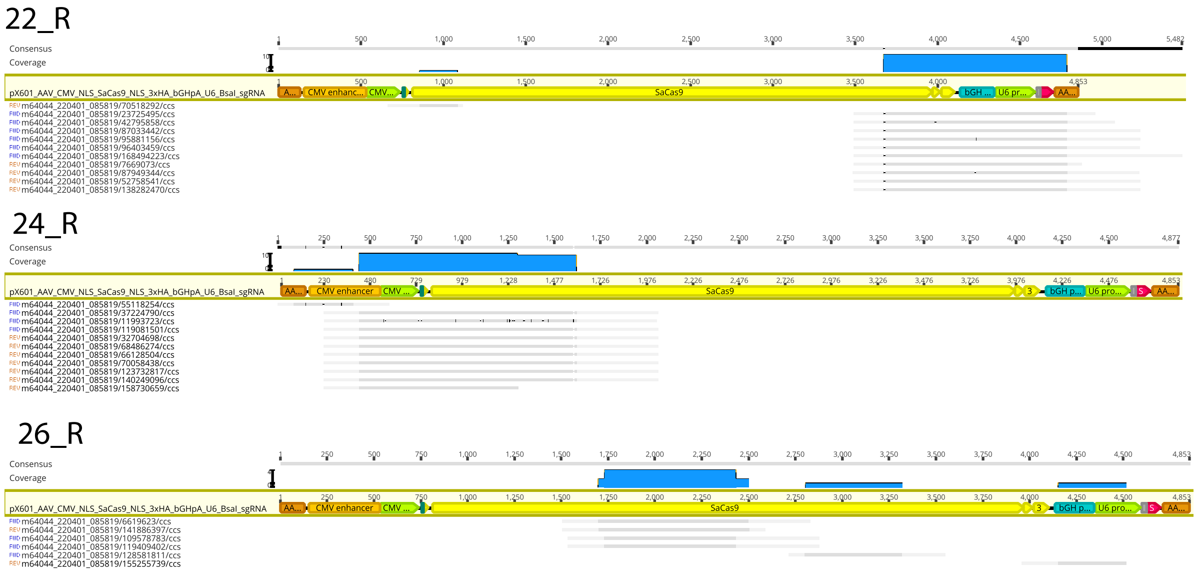


B

| Sample | AAV2 ITR 5Prime | CMV Enhancer | CMV Promoter | SV40 NLS | SaCas9 | Nucleoplasmin NLS | 3xHA | bGH polyA signal | U6 promoter | gRNA | AAV2 ITR 3Prime | Total_Reads_Aligned | Sequencing_Depth | Integration_Frequency (%) |
|---|---|---|---|---|---|---|---|---|---|---|---|---|---|---|
| 16_L (Sham) | 0 | 0 | 0 | 0 | 0 | 0 | 0 | 0 | 0 | 0 | 0 | 0 | 121273 | 0.0000 |
| 17_R (Sham) | 0 | 0 | 0 | 0 | 0 | 0 | 0 | 0 | 0 | 0 | 0 | 0 | 89472 | 0.0000 |
| 18_L (Sham) | 0 | 0 | 0 | 0 | 0 | 0 | 0 | 0 | 0 | 0 | 0 | 0 | 98593 | 0.0000 |
| 18_R (Sham) | 0 | 0 | 0 | 0 | 0 | 0 | 0 | 0 | 0 | 0 | 0 | 0 | 73703 | 0.0000 |
| 22_L (Cas9 AAV) | 0 | 0 | 0 | 0 | 0 | 0 | 0 | 0 | 0 | 0 | 0 | 0 | 112003 | 0.0000 |
| 22_R (Cas9 AAV) | 0 | 0 | 0 | 0 | 11 | 10 | 10 | 10 | 10 | 10 | 10 | 11 | 126532 | 0.0087 |
| 24_R (Cas9 AAV) | 1 | 11 | 10 | 10 | 10 | 0 | 0 | 0 | 0 | 0 | 0 | 11 | 109622 | 0.0100 |
| 25_R (Cas9 AAV) | 0 | 0 | 0 | 0 | 0 | 0 | 0 | 0 | 0 | 0 | 0 | 0 | 124005 | 0.0000 |
| 26_L (Cas9 AAV) | 0 | 0 | 0 | 0 | 0 | 0 | 0 | 0 | 0 | 0 | 0 | 0 | 119651 | 0.0000 |
| 26_R (Cas9 AAV) | 0 | 0 | 0 | 0 | 5 | 0 | 0 | 1 | 1 | 0 | 0 | 6 | 124038 | 0.0048 |
| 27_R (Cas9 AAV) | 0 | 0 | 0 | 0 | 0 | 0 | 0 | 0 | 0 | 0 | 0 | 0 | 131062 | 0.0000 |
| 31_R (Sham) | 0 | 0 | 0 | 0 | 0 | 0 | 0 | 0 | 0 | 0 | 0 | 0 | 115193 | 0.0000 |
| 32_L (Sham) | 0 | 0 | 0 | 0 | 0 | 0 | 0 | 0 | 0 | 0 | 0 | 0 | 132274 | 0.0000 |
| 32_R (Sham) | 0 | 0 | 0 | 0 | 0 | 0 | 0 | 0 | 0 | 0 | 0 | 0 | 113833 | 0.0000 |

C

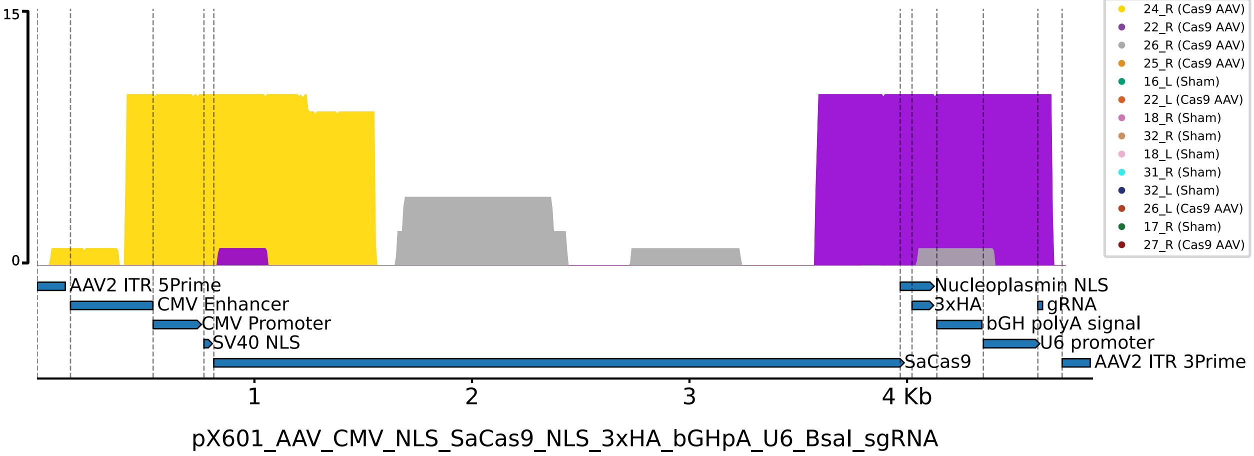


**Supplemental Figure 11. Long-read sequencing to detect viral fragment integrations at tdTomato locus following genome editing. A) Screen captures from Geneious software showing reads aligned to AAV reference genome (dark gray) with endogenous flanking regions (light gray) indicating viral genome trapping at tdTomato locus.** B) Tabulated reads from PacBio CCS within tdTomato locus that mapped to AAV genome components. The frequency of integration was low (< 0.01%), however estimated on target editing by ddPCR was <10% of alleles (Supplemental 8D). C) Visualization of reads mapping against the AAV genome. 3/7 sequenced brain samples that were injected with Cas9-AAV9 had detectable integration events, while 0/7 sequenced sham samples had detectable integrations.

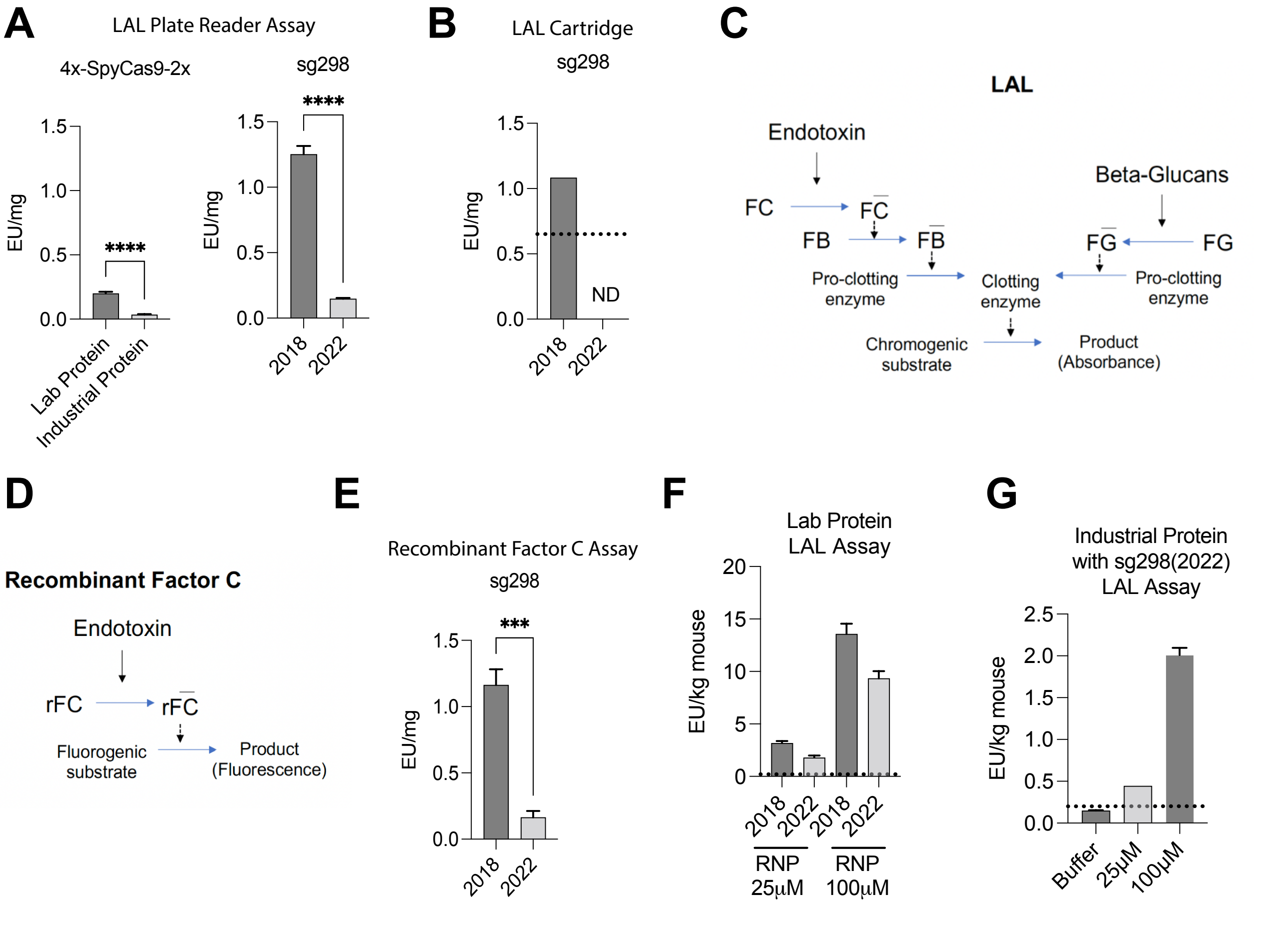


**Supplemental Figure 12. LAL and Recombinant Factor C assays to quantify endotoxins in RNPs.**

A) Limulus amebocyte lysate (LAL) plate reader based assay (Endosafe, Charles River) was performed on two lots of 4x-SpyCas9-2x protein and two lots of sg298, compared to a control standard endotoxin (CSE) from E.coli 055:B5 to generate endotoxin units (EU) per miligram (mg). Samples were run in triplicate and analyzed with unpaired t-tests,
**** p<0.0001.
B) LAL assay was repeated using the cartridge based system (Endosafe nexgen-PTS, Charles River), which reports a single value from two replicates with and without CSE spike-in internal controls. Statistical test was not performed. ND = not detected, dotted line represents limit of detection for the cartridge.
C-D) Schematic demonstrating the limulus amebocyte lysate (LAL) and recombinant factor C assays. The Recombinant Factor C assay has a fluorescent read out, negates the need for horsehsoe crab blood, and is not impacted by beta-glucans.
E) Recombinant Factor C assay (Pyrogene, Lonza) was performed on two lots of sg298 compared to a control standard endotoxin (CSE) from E.coli 055:B5 to generate endotoxin units (EU) per miligram (mg). Samples were run in triplicate and analyzed with unpaired t-tests, *** p<0.001.
F) Final RNP formulation (laboratory 4x-SpyCas9-2x protein with sg298 lots) had significantly more endotoxin than the FDA recommendation of 0.2 EU/kg/hr (dotted line) for human drug products administered into the central nervous system when delivered at 10µL in a 22g mouse.
G) The optimized RNP formulation (industrial 4x-SpyCas9-2x protein with sg298 from 2022) nearly reached the FDA recommendation of 0.2 EU/kg/hr (dotted line) at the 125pmol dose (25µM). Given the linear increase between 25-100µM the 10µM dose (50pmol) may be sufficient to reduce endotoxin below the threshold, while maintaining high levels of editing.

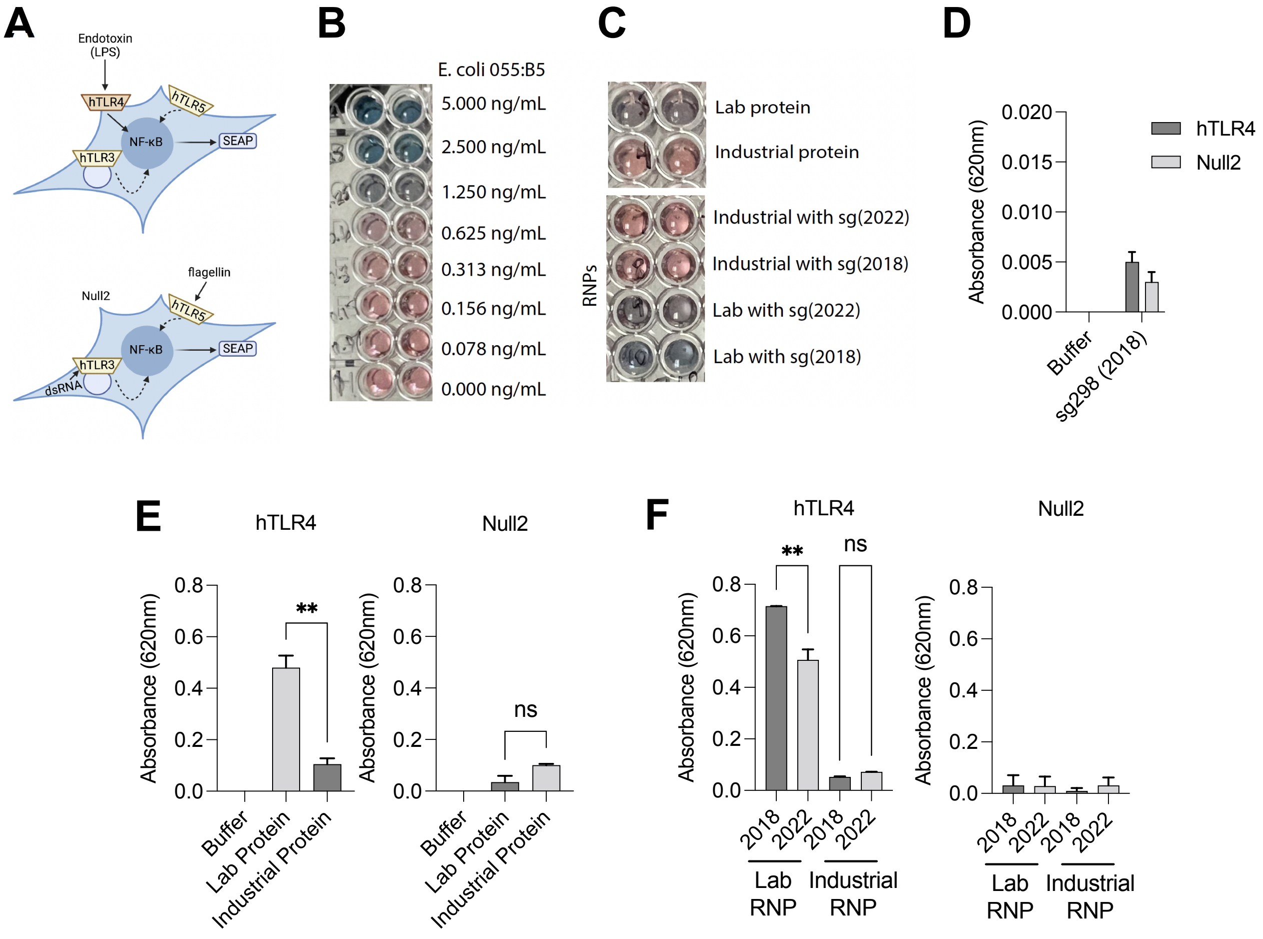


**Supplementary Figure 13. The effect of endotoxin in RNPs on NF-kB activation measured via HEK-Blue assay.**

A) HEK-Blue Assay (InvivoGen). Stimultaion with a TLR4 ligand, such as endotoxin / LPS, activates NFkappaB and AP-1, which induces secreted embryonic alkaline phosphotase (SEAP). SEAP can be easily detected in the cell-culture media and read with a plate reader. The parental cell line, Null2, will produceSEAP in response to TLR3, TLR5, and NOD1 agonists and is used to control for TLR4-specific activation of NFkappaB/AP-1.
B) Representative images of blue media change due to SEAP release downstream of NFkappaB/AP-1 activation following stimulation with control standard endotoxin.
C) Representative images of blue media change due to SEAP release downstream of NFkappaB/AP-1 activation following treatment with proteins and RNPs.
D) HEK-Blue assay applied to sg298 (2018) lot showed minimal change in absorbance from buffer. Assay was run in duplicate. Cells treated with sg298 lot from 2022 alone was not included in this experiment.
E) HEK-Blue Assay applied to proteins made in the lab or industrial scale-up. The lab-produced protein induced significantly more SEAP than the industrial protein. Assay was run in duplicate, unpaired t-tests, ** $p<0.01$.
F) HEK-Blue Assay applied to RNPs made with lab or industrial protein and sg298 from the 2018 or 2022 lots. The lab protein with the sg298 ("standard" formulation) was the most potent at inducing SEAP, while cells treated with the the industrial RNPs had much less SEAP production. Assay was run in duplicate, unpaired t-tests, ** $p<0.01$.

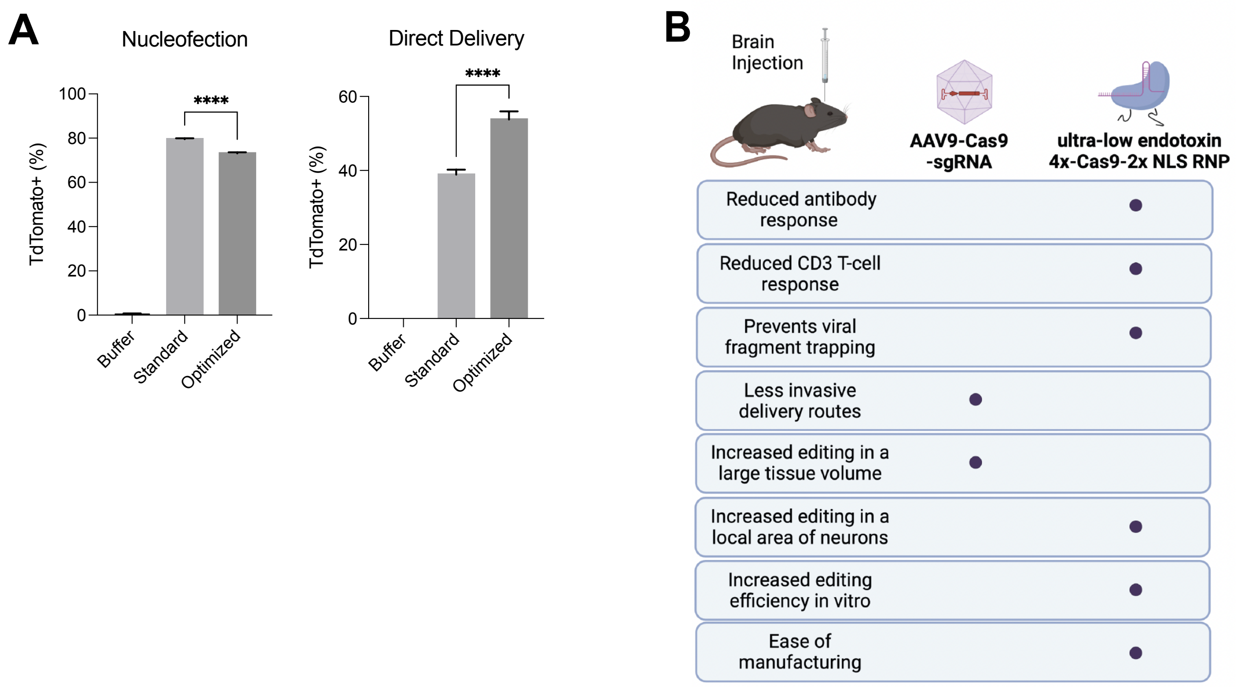


**Supplemental Figure 14. In vitro testing of optimized RNPs and final summary**.

A) Optimized RNPs (ultra-low endotoxin industrial protein with sg298 from 2020 lot) edited fewer Ai9 NPCs via nucleofection and more NPCs via direct delivery compared to the same dose (100pmol) of standard RNPs. This result showed similar trends to increasing the number of NLS on the N-terminus of SauCas9, suggesting more cell-penetrating peptides improve self-delivery / endosomal escape and reduce nucleofection efficacy (likely due to positive charge mechanism). These differences could also explain in-part the measured difference with editing in vivo (Figure 3), although the optimized RNPs were at least as efficacious in vivo as the standard formulation (Figure 1), demonstrating potential lot-to-lot variability in potency (active site concentration). Protein concentrations were confirmed via nanodrop before both in vitro and in vivo experiments.

B) Comparison of AAV9 and 4x-NLS RNP mediated delivery of Cas9 in the mouse central nervous system. To enable high-levels of editing in neurons within a localized brain region, minimize adaptive immune responses, and timely and affordable manufacturing scale up, the RNP offers an effective alternative delivery system to viral vectors. Image made using Biorender.com.
